# Differential propagation of ripples along the proximo-distal and septo-temporal axes of dorsal CA1 of rats

**DOI:** 10.1101/859082

**Authors:** Mekhala Kumar, Sachin S. Deshmukh

## Abstract

The functional connectivity of the hippocampus with its primary cortical input, the entorhinal cortex, is organized topographically. In area CA1 of the hippocampus, this leads to different functional gradients along the proximo-distal and septo-temporal axes of spatial/sensory responsivity and spatial resolution respectively. CA1 ripples, a network phenomenon, allows us to test whether the hippocampal neural network shows corresponding gradients in functional connectivity along the two axes. We studied the occurrence and propagation of ripples across the entire proximo-distal axis along with a comparable spatial range of the septo-temporal axis of dorsal CA1. We observed that ripples could occur at any location, but their probability of co-occurrence and amplitude decreased with increasing distance from the reference tetrode. This reduction was greater along the proximo-distal axis than the septo-temporal axis. Furthermore, we found that ripples propagate primarily along the proximo-distal axis. Thus, over a spatial scale of ~1.5 mm, the network is anisotropic along the two axes, complementing the topographically organized cortico-hippocampal connections.

## Introduction

The hippocampal proximo-distal and septo-temporal axes have an interesting connectivity pattern with their input regions. The hippocampus receives its primary cortical input from the entorhinal cortex (EC). The hippocampus and EC have a reciprocal anatomical connectivity showing two orthogonal patterns (Witter and Amaral, 2004). The medial entorhinal cortex (MEC) projects preferentially to proximal CA1, and the lateral entorhinal cortex (LEC) to distal CA1 (Steward and Scoville, 1976; Naber et al., 2001). Dorso-caudal MEC and lateral LEC project to septal hippocampus, while rostro-ventral MEC and medial LEC project to temporal hippocampus (Naber et al., 2001). Connections from CA1 and the subiculum to EC also maintain this topographical organization (Naber et al., 2001; Witter and Amaral, 2004), creating a loop. In addition to the differential connectivity along the two orthogonal axes of EC, the nature of information sent to the hippocampus by EC also varies along these axes. MEC shows path integration derived spatial representation, while LEC shows sensory derived non-spatial as well as spatial representation (Hafting et al., 2005; Deshmukh and Knierim, 2011). Grid spacing increases in discrete steps along the dorso-ventral axis of MEC (Stensola et al., 2012), though no such functional gradient has been demonstrated yet along the latero-medial axis of LEC. Corresponding to the functional differences in EC, the hippocampus shows functional differences along its two orthogonal axes. The proximo-distal axis of CA1 shows differences in spatial and sensory encoding (Henriksen et al., 2010; Oliva et al., 2016b), while the septo-temporal axis shows an increase in the scale of spatial representation (Jung et al., 1994; Maurer et al, 2005). In order to better understand the link between the structure and the function of the hippocampus described above, it is essential to understand how information flows through these networks involving the hippocampus.

The sharp-wave ripple complex is one such network phenomenon that may inform us about the hippocampal network organization. Depolarization of the apical dendrites of CA1 pyramidal cells by synchronous bursting of CA3 pyramidal cells causes sharp-waves in CA1. Ripples, high frequency oscillations (150-250 Hz) caused by the coordinated activation of CA1 pyramidal cells and interneurons, often accompany sharp-waves. These sharp-wave ripple complexes are observed during periods of awake immobility, non-REM sleep, and consummatory behaviors (Buzsàki et al., 1992; Ylinen et al., 1995; Csicsvari et al., 1999; Buzsàki, 2015).

Along the septo-temporal axis, ripples show a reduction in amplitude as a function of distance from any reference electrode, and a tendency to propagate (Patel et al., 2013). We tested if these patterns hold along the proximo-distal axis. We compared these proximo-distal patterns with those along a comparable spatial extent along the septo-temporal axis in dorsal CA1 in our dataset. We show that there is a decrease in amplitude and probability of co-occurrence as a function of distance from the reference, and that ripples travel preferentially along the proximo-distal axis.

## METHODS

### Animals and Surgery

All animal procedures were performed in accordance with the National Institutes of Health (NIH) animal use guidelines and were approved by the Johns Hopkins University Animal Care and Use Committee. Local field potentials (LFPs) were recorded from eight male, 5-6 month old Long Evans rats. A hyperdrive was implanted over the right dorsal hippocampus (see Deshmukh and Knierim, 2011 for details of surgical procedures). The hyperdrive had ~3 rows of guide cannulae (15 independently movable tetrodes and 2 references) angled at 35° to the ML axis and the lateral/anterior-most tetrode was positioned at ~3.4 mm posterior and 3.4 mm lateral to bregma in order to record from the entire proximo-distal axis of CA1.

### Behavioral Protocol

All rats were trained and tested on a circular track (see Knierim, 2002 for details). Resting sessions were recorded while the rats were made to sit on a pedestal for about 20 minutes before the first session and after the last session on each day. LFPs from only the resting sessions were analyzed and have been presented here. We neither recorded a video nor kept track of the behavioral state of the rats during these resting sessions. Thus, we could not segregate the periods of awake immobility from the periods of sleep.

### Data Acquisition

Neural recordings were performed using a 64 channel wireless transmitter (Triangle Biosystems, Durham, NC) connected to a Cheetah Data Acquisition system (Neuralynx, Bozeman, MT). LFPs were recorded from one channel from each tetrode after amplifying 2000-fold, filtering between 1 and 475 Hz, and digitizing at 1 kHz.

### Histology

Histological reconstruction was used to localize tetrodes during each recording session. For quantitative purposes, relative distances along the proximo-distal and septo-temporal axes were estimated from histology.

### Ripple detection

Theta epochs were excluded from the analysis. To detect ripples, the root mean square (RMS) amplitude of LFPs filtered between 150-250 Hz was calculated using a moving window of 10 ms. A threshold of 2 SD above mean was then used to first detect putative ripple events. The start and end time points (boundaries) of these putative events were detected using a threshold of 1 SD above mean, and only the events with durations between 25 ms and 200 ms were classified as ripples. In addition to collectively analyzing all ripple events ≥ mean + 2 SD, the ripple events were classified into amplitude groups of mean + 2-4 SD, mean + 4-6 SD, and ≥ mean + 6 SD and analyzed separately. These will be referred to as ≥ 2 SD, 2-4 SD, 4-6 SD, and ≥ 6SD through the rest of the manuscript. The time of the peak amplitude of a ripple obtained from the envelope of the analytical signal was used as the time of occurrence of the ripple.

In the analyses that follow, each tetrode served as a reference against which every other tetrode was compared.

### Data considered for analysis

LFP recordings from only the tetrodes in the pyramidal cell layer of CA1 showing ripples were included (Figure 1). For each rat, a total of 4-8 resting sessions were pooled. The rate of ≥ 2SD ripple occurrence across sessions and rats was 47 ± 10 ripples/min. In all, the dataset included 305548 ripples from 58 tetrodes in 38 sessions in 8 rats.

**Figure 1.**
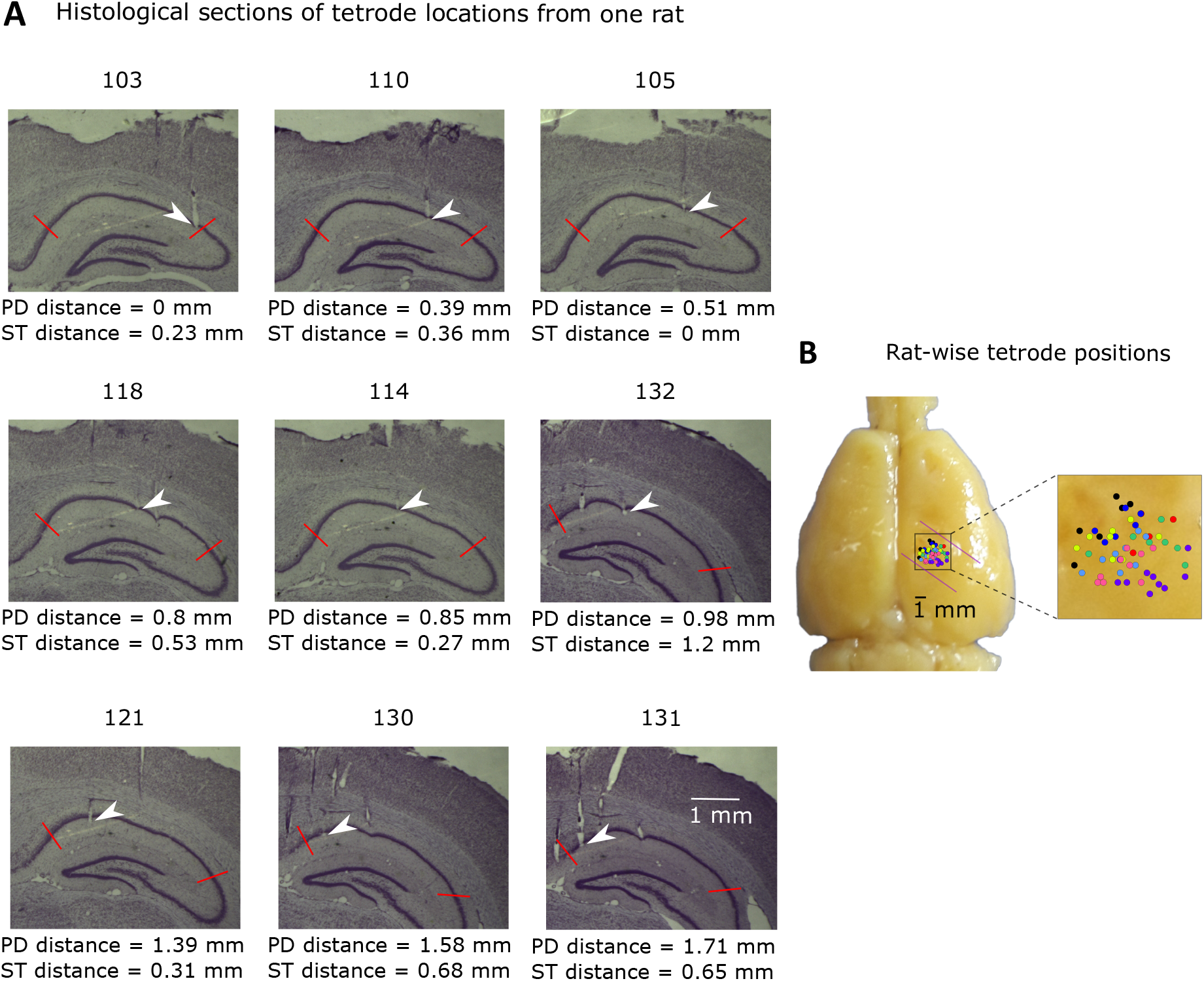
Tetrode positions. (A) Histological sections showing tetrode locations from one rat (rat 441). Arrows indicate tetrodes included in the analysis. Other tracks belong to tetrodes excluded from the analysis since they were not in the CA1 pyramidal cell layer on the days included in the analysis. Numbers atop each section indicate the section number. Sections were 47 μm thick after accounting for 15% histological shrinkage factor. (B) Distribution of all tetrodes used for the analysis across the proximo-distal and septo-temporal axes of CA1. Dots of a color represent tetrodes from a single rat. Relative tetrode positions obtained from histology were marked on the whole brain for display purposes using the estimates obtained from the rat brain atlas (Paxinos and Watson, 2007). Purple lines indicate the orientation of the septo-temporal axis.

### Detection of co-occurring ripples

A ripple on a referred tetrode was considered to co-occur with a ripple on a reference tetrode if there was an overlap between the boundaries of the two ripples. To reduce the likelihood of a ripple co-occurring on the referred tetrode going undetected due to a reduction in amplitude, the threshold for ripple detection on the referred tetrode was maintained at 2 SD above mean (other criteria remained unchanged) irrespective of the threshold on the reference tetrode. For each reference-referred pair, the fraction of ripples on the reference tetrode having co-occurring ripples on the referred tetrode was computed.

### Measurement of relative amplitude

The amplitude of a ripple was measured as the peak with the largest amplitude in an identified ripple. If the signal on the referred tetrode was below the ripple detection threshold, the peak amplitude on the referred tetrode was measured within the boundaries of the ripple on the reference tetrode. A variety of factors, including exact location of the tetrode in the pyramidal cell layer (Ylinen et al., 1995; Mizuseki et al., 2011) as well as tetrode impedance affect the recorded ripple amplitude. To account for tetrode to tetrode variability, amplitudes measured on each tetrode were normalized by the median ripple amplitude on that tetrode. The ratio of the normalized amplitude on the referred tetrode to the normalized amplitude on the reference tetrode was calculated as the relative amplitude. For each reference-referred pair, the mean of all the relative amplitudes was computed.

### Propagation of ripples

In order to understand ripple propagation, each of the four extreme tetrodes (the proximal-most, distal-most, septal-most, and temporal-most tetrodes) in each of the hyperdrives were used as reference tetrodes to detect ripples. Ripples were then sequentially detected on the referred tetrodes in order of increasing distance from the reference tetrode overlapping with the co-occurring ripple on the preceding tetrode to identify a putative propagating event. Any common events detected between the four extreme reference tetrodes were counted only once. Only those events having at least six consecutive tetrodes with ripples and spanning a minimum distance of 0.6 mm along the proximo-distal and septo-temporal axes were analyzed further (rat 305 had only 4 tetrodes and was, therefore, excluded). Multiple linear regression was then performed on the time of occurrence of ripples vs. relative distance along the proximo-distal and septo-temporal axes. An event was considered as propagating if the fit was significant (at p < 0.05). Some events were common between the 2-4 SD, 4-6 SD, and ≥ 6 SD amplitude groups, e.g. if an event having a ripple falling in the amplitude group of ≥ 6 SD on one extreme tetrode had a ripple falling in the amplitude group of 2-4 SD on another extreme tetrode, the event would be detected for 2-4 SD as well as ≥ 6 SD. These events were counted only once by including the events only in the highest amplitude group that they belonged to.

To find the direction of propagation of each of these events, vector algebra was employed. The partial slopes along the two orthogonal axes viz. the proximo-distal and septo-temporal axes obtained from the multiple linear regression analysis were considered as component vectors, their magnitude defined by the absolute value of the partial slope and their direction along their respective axis defined by the sign (+/−) of the partial slope. The resultant of the two orthogonal component vectors was then computed to obtain the resultant slope and angle of propagation, which was calculated with respect to the proximo-distal direction. Thus, propagation along the proximo-distal and disto-proximal directions were defined by the angles 0°, and 180° or −180° respectively, while propagation along the septo-temporal and temporo-septal directions were defined by the angles 90° and −90° respectively. Intermediate directions were defined by intermediate angles.

The average speed along each of the four directions was measured by first categorizing events into the four directions of propagation using bin widths of ±45° about the expected angle of each of the four directions. For each direction, the relative time of occurrence of all ripples on each tetrode with respect to the ripples on the tetrode with the earliest time of occurrence was averaged across events. Multiple linear regression was performed on each of the four averaged datasets, and the average speed for each of the four directions was computed as the inverse of the resultant slope. Additionally, the speed was separately calculated for each propagating event as the inverse of the resultant slope for each event.

To ensure that propagation in the observed preferred direction and speed did not occur by chance, a shuffle analysis was performed. For each propagating event, the time of occurrence of each of the ripples was randomly permuted with respect to the tetrode positions to generate the shuffled dataset. The multiple linear regression analysis and significance criteria were then applied to each of the shuffled events as was done for the observed data. This entire process was done iteratively to generate 1000 such datasets; for a given rat, for every iteration a single random permutation of the time of occurrence with respect to tetrode positions was used for all propagating events. To quantify the possible differences in the preferred direction of propagation (proximo-distal vs. septo-temporal) between the observed data and shuffle data, we calculated the ratio of the number of events propagating along the proximo-distal and disto-proximal directions to those along the septo-temporal and temporo-septal directions:

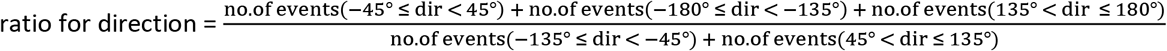

where “dir” denotes direction. In addition to this, for the observed data, the distribution of the partial slopes along the proximo-distal axis was bimodal but that along the septo-temporal axis was not.

Therefore, we calculated the following to quantify possible differences in the distribution of the partial slopes of the observed and shuffled data:

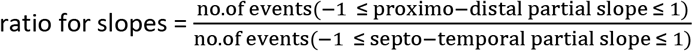

A range of ± 1 ms/mm was used as the separation in the two populations along the proximo-distal axis was evident within this range for the observed data. The observed data was considered to be significantly different from chance if less than 1% of the 1000 shuffled iterations had ratios for direction more than that of the observed data and ratios for slopes less than that of the observed data.

### Statistical analysis

All statistics used a significance criterion of p < 0.05 unless mentioned otherwise. One-tailed t-tests were used to compare partial slopes along the proximo-distal and septo-temporal axes.

## RESULTS

We recorded LFPs across the entire proximo-distal axis and a comparable spatial range along the septo-temporal axis from the dorsal CA1 of rats (Figure 1) during resting sessions. Ripples were detected using a threshold of 2 SD above mean. All ripples ≥ 2 SD were collectively analyzed, and additionally, were classified into 2-4 SD, 4-6 SD, and ≥ 6 SD and analyzed separately. To look for possible differences in the occurrence of ripples, we first examined whether the rate of ripple occurrence was influenced by the positions of the tetrodes along the proximo-distal and septo-temporal axes. We observed no differences in the rates for ≥ 2 SD (Figure 2A; proximo-distal partial slope = −1.2 count/min/mm, p = 0.54; septo-temporal partial slope = −1.5 count/min/mm, p = 0.59; one-tailed t-test for comparison of slopes along the two axes, p = 0.53). However, the rate was higher in proximal CA1 than distal CA1 for 2-4 SD (Figure 2B), did not show a significant trend for 4-6 SD (Figure 2C), and was higher in distal CA1 than proximal CA1 for ≥ 6 SD (Figure 2D). There was no such gradient along the septo-temporal axis for any of the three subgroups (Figure 2B-D; see Supplementary Table 1 for statistics after subdividing ripples by amplitude and Supplementary Table 2 for ripple rate as a function of tetrode position in individual rats). After controlling for differences in the spatial spreads of tetrodes across the proximo-distal and septo-temporal axes, there were no significant differences in ripple rates as a function of tetrode position (Supplementary Figure 1, Supplementary Table 3).

Next, we asked whether these ripples co-occurred with each other (Figure 3). For each reference tetrode, we looked at the fraction of ripples having co-occurring ripples on all the referred tetrodes. We quantified the correlation between co-occurrence and distance by plotting the fraction of co-occurring ripples as a function of the absolute distance between all pairs of tetrodes along both axes (Figure 4). The likelihood of co-occurrence decreased with increasing distance for ≥ 2 SD (Figure 4A). This decrease was significant along the proximo-distal axis but not the septo-temporal axis (proximo-distal partial slope = −0.071 fraction of co-occurring ripples/mm, p = 2 × 10-12*; septo-temporal partial slope = −0.019 fraction of co-occurring ripples/mm, p = 0.14) and was significantly steeper along the proximo-distal axis than the septo-temporal axis (one-tailed t-test for comparison of slopes along the two axes, p = 4 × 10^−4*^). Consistent with this, the fraction of co-occurring ripples decreased with increasing distance for 2-4 SD, 4-6 SD, and ≥ 6 SD (Figure 4B-D). Notice that the reference ripples in the lower amplitude group were less likely to have co-occurring ripples even for short distances between tetrode pairs, while reference ripples in the higher amplitude group were much more likely to have co-occurring ripples. The decrease in the fraction of co-occurring ripples was significant along the proximo-distal axis for all three subgroups. The decrease was not significant along the septo-temporal axis for 2-4 SD, but was significant for 4-6 SD and ≥ 6 SD. The rate of decrease in the fraction of co-occurring ripples was significantly higher along the proximo-distal axis than along the septo-temporal axis (see Supplementary Table 4 for statistics after subdividing ripples by amplitude on the reference tetrode and Supplementary Table 5 for ripple co-occurrence as a function of relative distance in individual rats). The results remained similar after controlling for the direction from the reference to the referred tetrode along the two axes (e.g. by segregating comparisons between pairs having more proximal and septal reference tetrodes than their corresponding referred tetrodes from pairs having more distal and temporal reference tetrodes than their referred tetrodes; Supplementary Figure 2, Supplementary Table 6) and differences in the spatial spreads of tetrodes across the proximo-distal and septo-temporal axes (Supplementary Figure 3, Supplementary Table 7).

**Figure 2.**
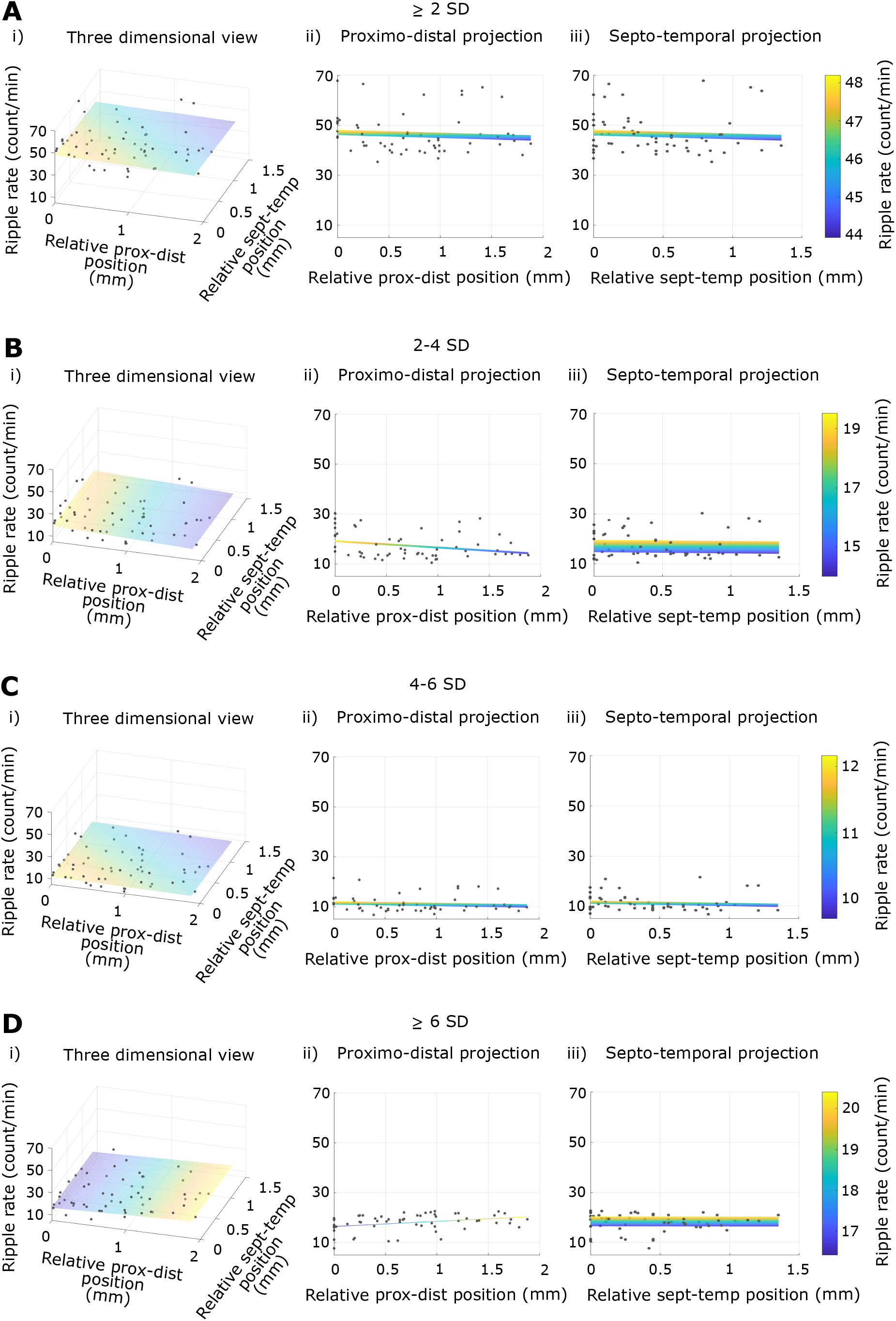
Rate of ripple occurrence. Rate of ripple occurrence as a function of position along the proximo-distal and septo-temporal axes for all rats collectively with the 2D fit (plane) for ≥ 2 SD (A), 2-4 SD (B), 4-6 SD (C), and ≥ 6 SD (D). Column (i)shows a 3D (X-Y-Z) view, (ii) shows a proximo-distal projection (X-Z view), and (iii) shows a septo-temporal projection (Y-Z view) of column (i). Note the decreasing trend in (B) and the increasing trend in (D) along the proximo-distal axis.

**Figure 3.**
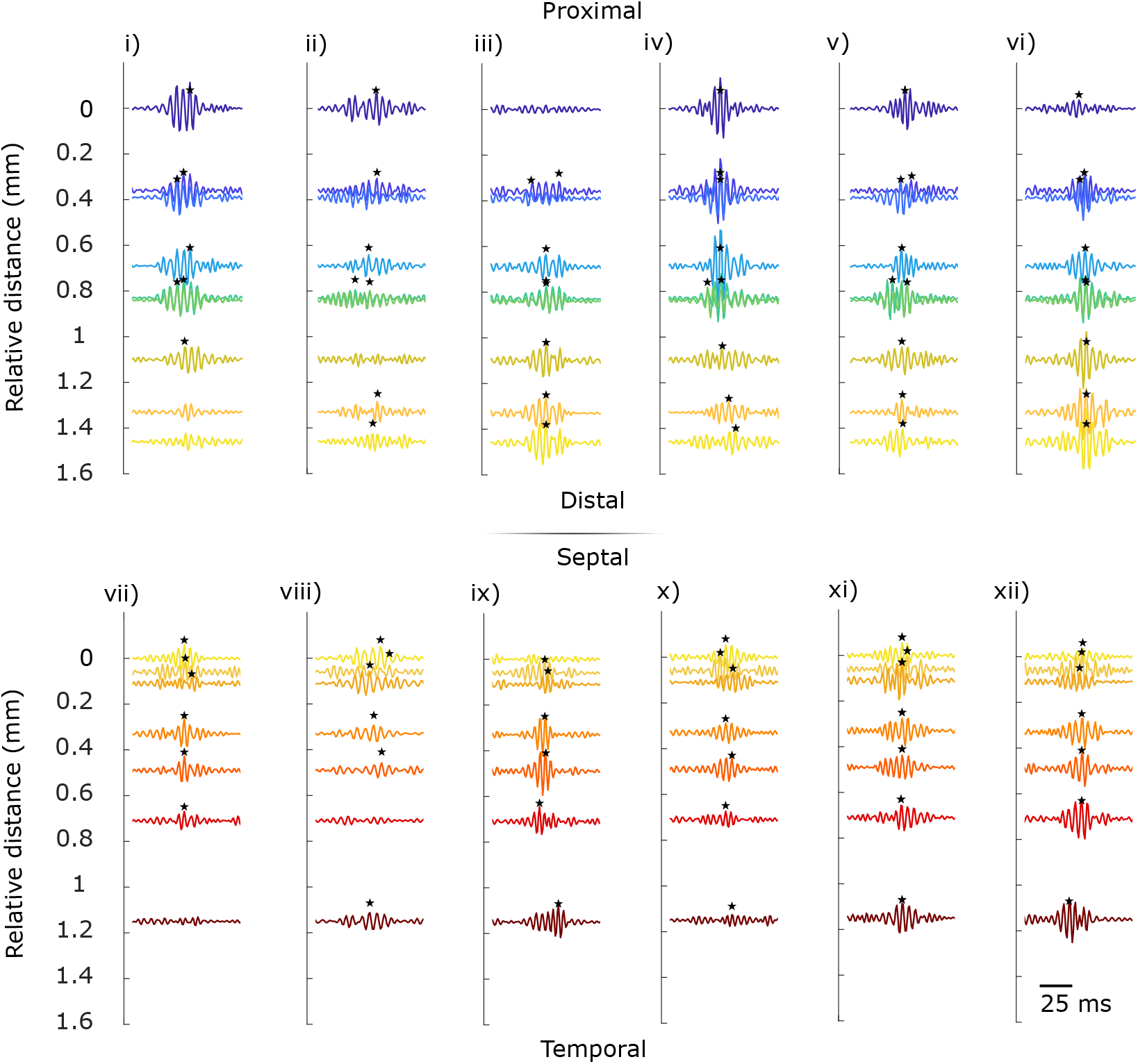
Examples of ripple patterns across tetrodes. Recorded traces are arranged from proximal to distal in (i)-(vi) (rat 432) while traces are arranged from septal to temporal in (vii)-(xii) (rat 392). Stars indicate detected ripples on a tetrode and are positioned at the time of peak amplitude of the ripples. (i)-(iii) and (vii)-(ix) show absence of co-occurring ripples on some of the tetrodes. (iv)-(vi) and (x)-(xii) show ripples of different amplitudes co-occurring on all tetrodes.

**Figure 4.**
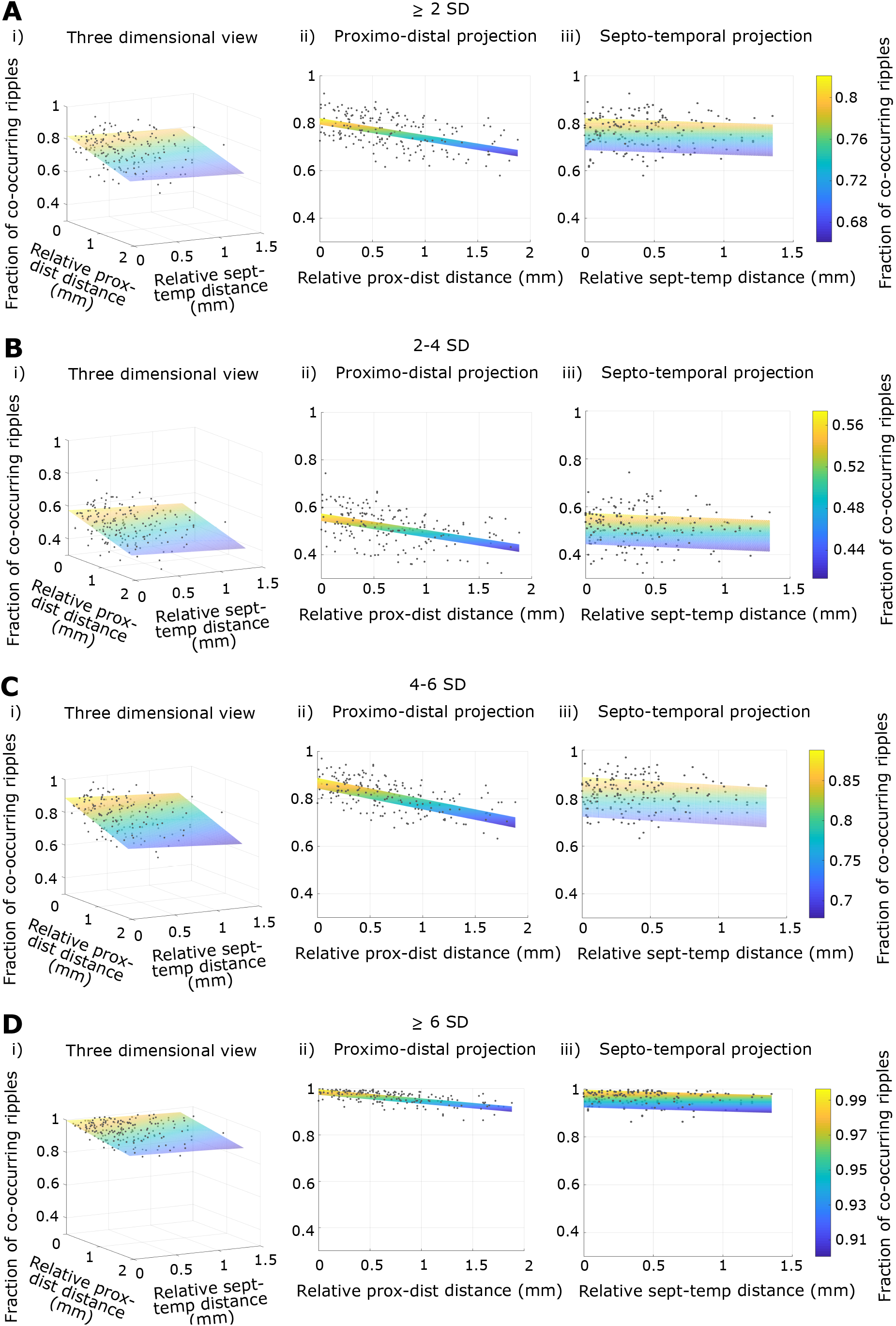
Ripple co-occurrence. Fraction of co-occurring ripples as a function of relative distance along the proximo-distal and septotemporal axes for all rats collectively with the 2D fit (plane) for ≥ 2 SD (A), 2-4 SD (B), 4-6 SD (C), and ≥ 6 SD (D) ripples on reference tetrodes. Column (i) shows a 3D (X-Y-Z) view, (ii) shows a proximo-distal projection (X-Z view), and (iii) shows a septo-temporal projection (Y-Z view) of column (i).

The decrease in co-occurrence with increasing distance suggested that there may be an amplitude gradient as a function of distance. To test the dependence of ripple amplitude on distance, we measured the relative amplitude of ripples on the referred tetrodes with respect to all ripples on the reference tetrode, irrespective of whether a co-occurring ripple meeting our criterion (mean + 2 SD) was detected on the referred tetrode. As expected, the relative amplitude of ripples decreased with increasing (absolute) distance for ≥ 2 SD; the decrease was significant along the proximo-distal axis but not the septo-temporal axis (Figure 5A; proximo-distal partial slope = −0.025 relative amplitude/mm, p = 2 × 10^−5*^; septo-temporal partial slope = −0.005 relative amplitude/mm, p = 0.56), with the decrease being significantly steeper along the proximo-distal axis than the septo-temporal axis (one-tailed t-test for comparison of slopes along the two axes, p = 0.013*). For 2-4 SD and 4-6 SD, there was neither an increase nor a decrease along either axis (Figure 5B,C), but ≥ 6 SD showed a significant decrease in relative amplitude along both axes with the decrease being steeper along the proximo-distal axis (Figure 5D; see Supplementary Table 8 for statistics after subdividing ripples by amplitude on the reference tetrode and Supplementary Table 9 for relative ripple amplitude as a function of relative distance in individual rats). The results remained similar after controlling for the direction from the reference to the referred tetrode along the two axes, except the decrease was no longer significant for ≥ 2 SD (Supplementary Figure 4, Supplementary Table 10). The results also remained similar after controlling for the differences in the spatial spreads of tetrodes across the proximo-distal and septo-temporal axes, except the decrease was no longer significant for ≥ 2 SD and no longer significantly steeper along the proximo-distal axis than the septo-temporal axis for ≥ 6 SD (Supplementary Figure 5, Supplementary Table 11).

**Figure 5.**
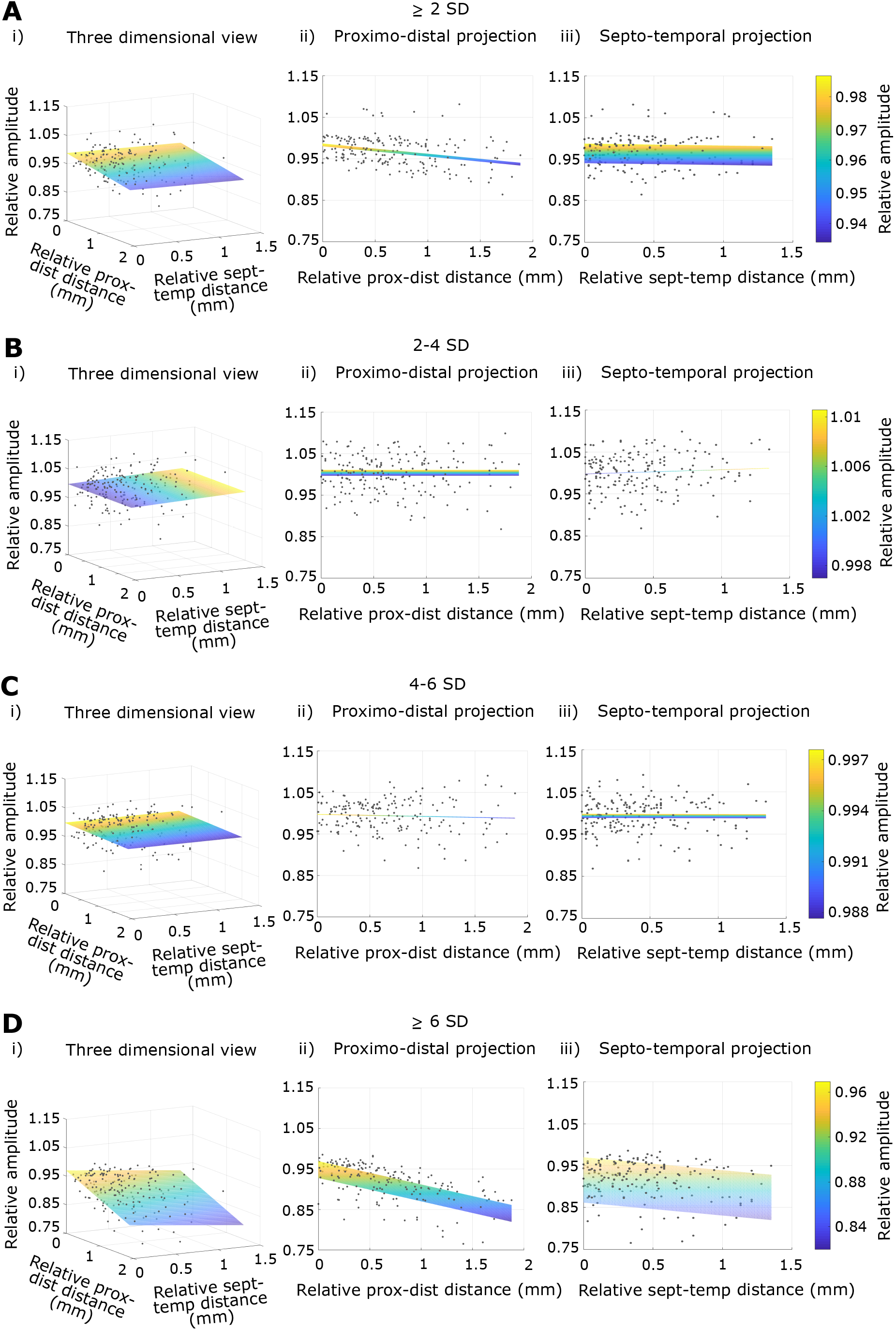
Relative ripple amplitude. Figure organization is the same as Figure 4, but for relative ripple amplitude. Note the absence of gradients along the proximo-distal and septo-temporal axes in (B) and (C).

Ripples have been shown to propagate over a few mm along the septo-temporal axis (Patel et al., 2013). We tested if ripples propagate similarly along the proximo-distal axis, as well as a matching spatial scale (~1.5 mm) along the septo-temporal axis (Figure 6). For ≥ 2 SD, out of the 25282 putative propagating events analyzed, 5199 events (20%) showed significant propagation (at p <0.05 for 2D fit). The distribution of the angles of propagation (with respect to the proximo-distal axis) of these events demonstrated a clear preference around 0° (1725 events within ±45° out of a total of 5199 propagating events, 33%) and 180° and −180° (1559 events, 30%) over propagation around −90° (972 events, 19%) and 90° (943 events, 18%) (Figure 7A; ratio for direction = 1.76; test of proportions comparing the observed proportion of events with the 50% propagation along the proximo-distal axis expected by chance, Z = 20, p = 0*) implying that the proximo-distal axis was the preferred axis of propagation. Additionally, the distribution of the proximo-distal partial slopes about 0 ms/mm was bimodal, unlike the septo-temporal partial slopes; there were a smaller number of events having proximo-distal partial slopes between ± 1 ms/mm as opposed to the greater number of events with septo-temporal partial slopes between ± 1 ms/mm (ratio = 0.346; Figure 7B). A slope close to 0 ms/mm corresponds to near simultaneous occurrence, and hence a lack of propagation along the given axis. In order to ensure that these results were not an outcome of the criteria used for classifying the events as propagating, we ran a shuffle analysis. The distributions of the angles of propagation for the shuffled data were significantly different from that of the observed data (ratio for direction for the observed data, 1.76, was higher than that for 994 shuffles, p = 0.006*), and so were the distributions of the partial slopes (ratio for slopes for the observed data, 0.346, was lower than 994 shuffles, p = 0.006*), thus indicating that the observed result was not by chance. Subdividing putative propagating events by amplitude on the reference tetrode revealed that the bias for propagation along the proximo-distal axis was higher for subgroups with higher amplitude on the reference tetrode (Figure 7C; Supplementary Figures 6,7,8; Supplementary Table 12).

**Figure 6.**
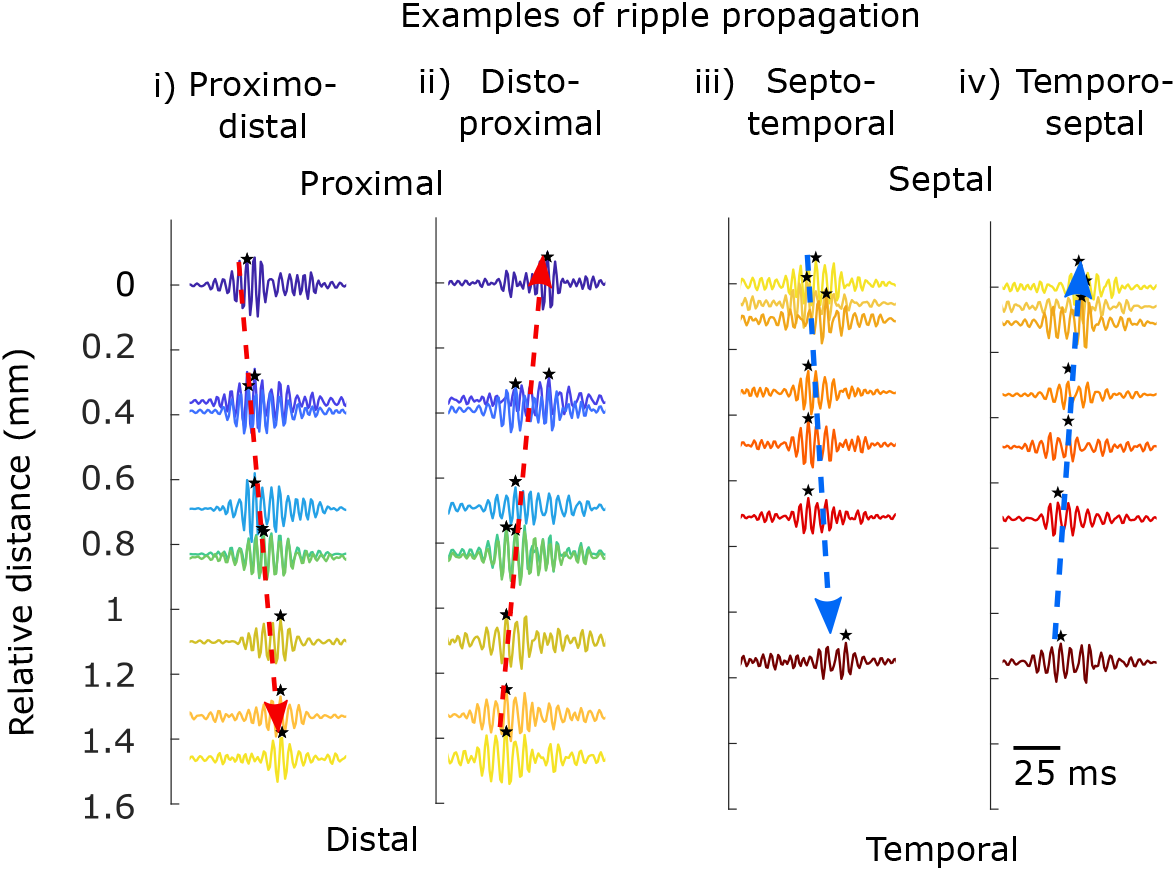
Examples of ripple propagation. Examples of ripples propagating from proximal to distal (rat 432) (i), distal to proximal (rat 432) (ii), septal to temporal (rat 392) (iii), and temporal to septal (rat 392) (iv). Arrangement of tetrodes is the same as in Figure 3. Dotted arrows indicate the direction of propagation.

**Figure 7.**
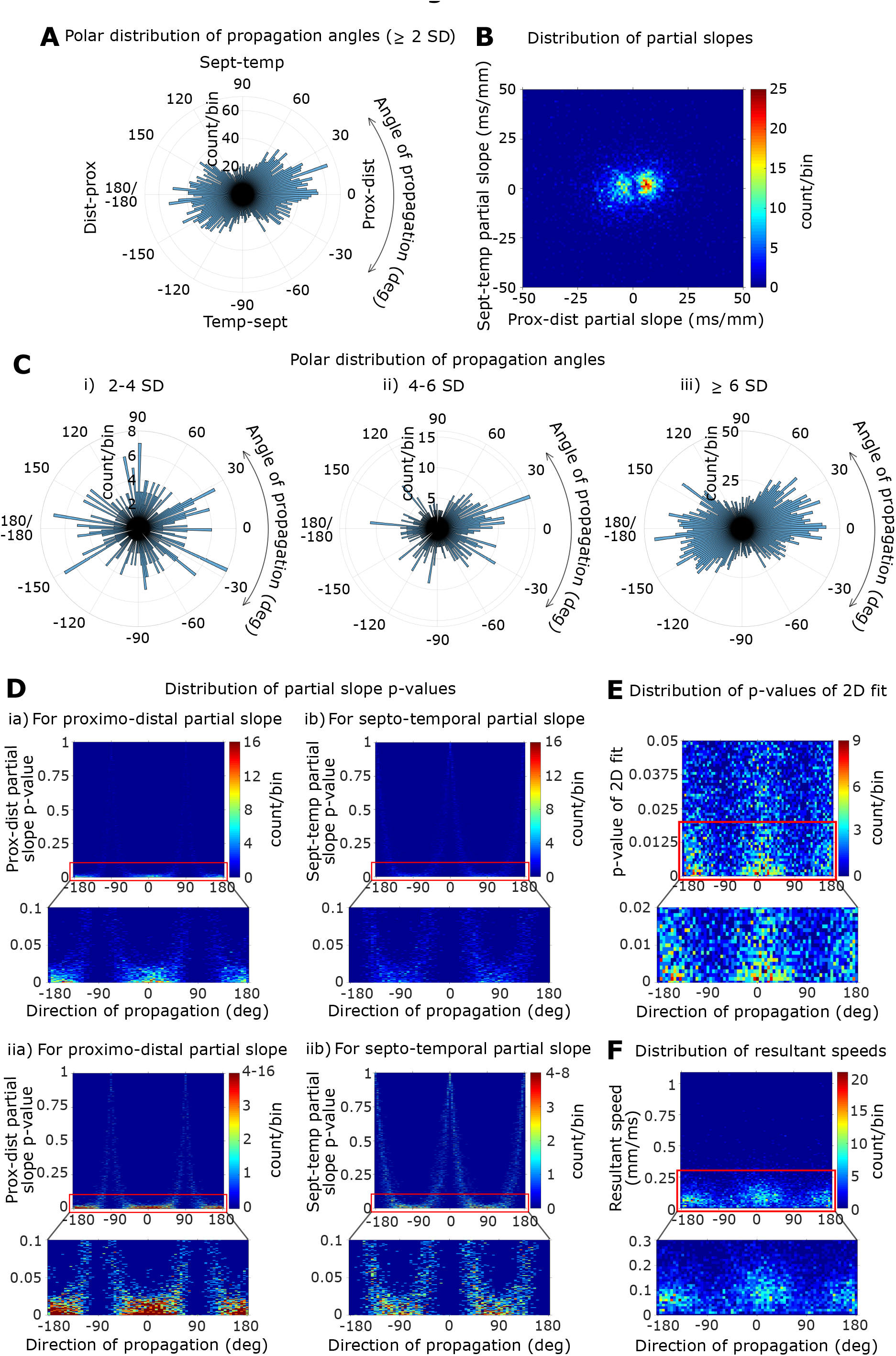
Ripple propagation. (A) Polar distribution of the angles of propagation for ≥ 2 SD. The following angles correspond to the given directions: proximo-distal direction, 0°; disto-proximal direction, 180° or −180°; septo-temporal direction, 90°; temporo-septal direction, −90°. Note the clear preference for the proximo-distal axis over the septo-temporal axis. (B) Distribution of proximo-distal vs. septo-temporal partial slopes obtained from multiple linear regression analysis of individual propagating events (≥ 2 SD). Note the bimodal nature of the distribution of the proximo-distal partial slopes but not the septo-temporal partial slopes. (C) Polar distribution of the angles of propagation for 2-4 SD (i), 4-6 SD (ii), and ≥ 6 SD (iii). (D) Distributions of the p-values of partial slopes vs. direction (angle) of propagation for all propagating events (≥ 2 SD). Panels (i) and (ii) are the same plots, however, the color scheme of (ii) has an upper limit of 4 (instead of the actual maximum bin count) to facilitate visualization of the pattern – note that the p-values are close to 0 about 0°, −180°, and 180° and close to 1 about −90° and 90° in (iia), and vice versa for the p-values in (iib). Each figure shows a zoomed in section below. (E) Distribution of the p-values of the multiple linear regression models vs. direction of propagation for all propagating events. (F) Distribution of the resultant speed (obtained from vector analysis) vs. direction of propagation for all propagating events.

It is possible that events preferentially propagating along the proximo-distal axis – events having relatively large, significant proximo-distal partial slopes – also had relatively small but significant septo-temporal slopes and vice versa for those propagating along the septo-temporal axis. Most of the events preferentially propagating in the proximo-distal direction showed high p-values for septo-temporal partial slopes, indicating that these events did not propagate significantly along the septo-temporal axis. Similarly, events preferentially propagating along the septo-temporal axis showed high p-values for proximo-distal partial slopes (Figure 7D, Supplementary Figure 6,7,8), indicating that these events did not propagate significantly along the proximo-distal axis. For ≥ 2 SD, 4-6 SD, and ≥ 6 SD, notice the substantially higher density of low p-values along the proximo-distal axis than that along the septo-temporal axis. Furthermore, the distribution of the p-values of the multiple regression models for all events also showed a greater number of events with p-values close to 0 along the proximo-distal axis than that along the septo-temporal axis (Figure 7E, Supplementary Figure 7,8). The distribution of propagation speeds of individual events as a function of direction showed that most events propagate at speeds < 0.2 mm/ms, and for ≥ 2 SD, 4-6 SD, and ≥ 6 SD they are concentrated at directions closer to the proximo-distal axis (Figure 7F, Supplementary Figure 7,8). Together, these results indicate that ripples either propagate along the proximo-distal or septo-temporal axes, and preferentially propagate along the proximo-distal axis for events having higher amplitude ripples.

Next, we estimated average ripple propagation speeds. Events were classified based on the four directions of propagation (Figure 6) and analyzed separately. On average, each group showed a clear preference for the expected direction of propagation (Figure 8). For propagation along the proximo-distal axis, the proximo-distal partial slopes were significantly larger than the septo-temporal partial slopes (for ≥ 2 SD: proximo-distal propagation: proximo-distal partial slope = 12.3 ms/mm, p = 3 × 10^−33*^; septo-temporal partial slope = 0.22 ms/mm, p = 0.7; one-tailed t-test for comparison of slopes along the two axes, p = 0*; disto-proximal propagation: proximo-distal partial slope = −13.2 ms/mm, p = 1 × 10^−23*^; septo-temporal partial slope = −0.05 ms/mm, p = 0.96; one-tailed t-test for comparison of slopes along the two axes, p = 2 × 10^−14*^). For propagation along the septo-temporal axis, the septo-temporal partial slopes were significantly larger than the proximo-distal partial slopes (for ≥ 2 SD: septo-temporal propagation: proximo-distal partial slope = 1.33 ms/mm, p = 0.24; septo-temporal partial slope = 15.9 ms/mm, p = 2 × 10^−10*^; one-tailed t-test for comparison of slopes along the two axes, p = 8 × 10^−9*^; temporo-septal propagation: proximo-distal partial slope = 0.77 ms/mm, p = 0.45; septo-temporal partial slope = −17.9 ms/mm, p = 5 × 10^−12*^; one-tailed t-test for comparison of slopes along the two axes, p = 1 × 10^−10*^). The patterns remained the same for 2-4 SD, 4-6 SD and ≥ 6 SD (see Supplementary Table 12 for statistics after subdividing ripples by amplitude and Supplementary Table 13 for statistics of ripple propagation in individual rats). For ≥ 2 SD, ripples propagated at 0.081 mm/ms in the proximo-distal direction, 0.076 mm/ms in the disto-proximal direction, 0.063 mm/ms in the septo-temporal direction, and 0.056 mm/ms in the temporo-septal direction (see Supplementary Figure 9,10,11, and Supplementary Table 12 for propagation speeds after subdividing propagating events by amplitude on reference tetrode; see Supplementary Table 13 for ripple propagation statistics for individual rats).

**Figure 8.**
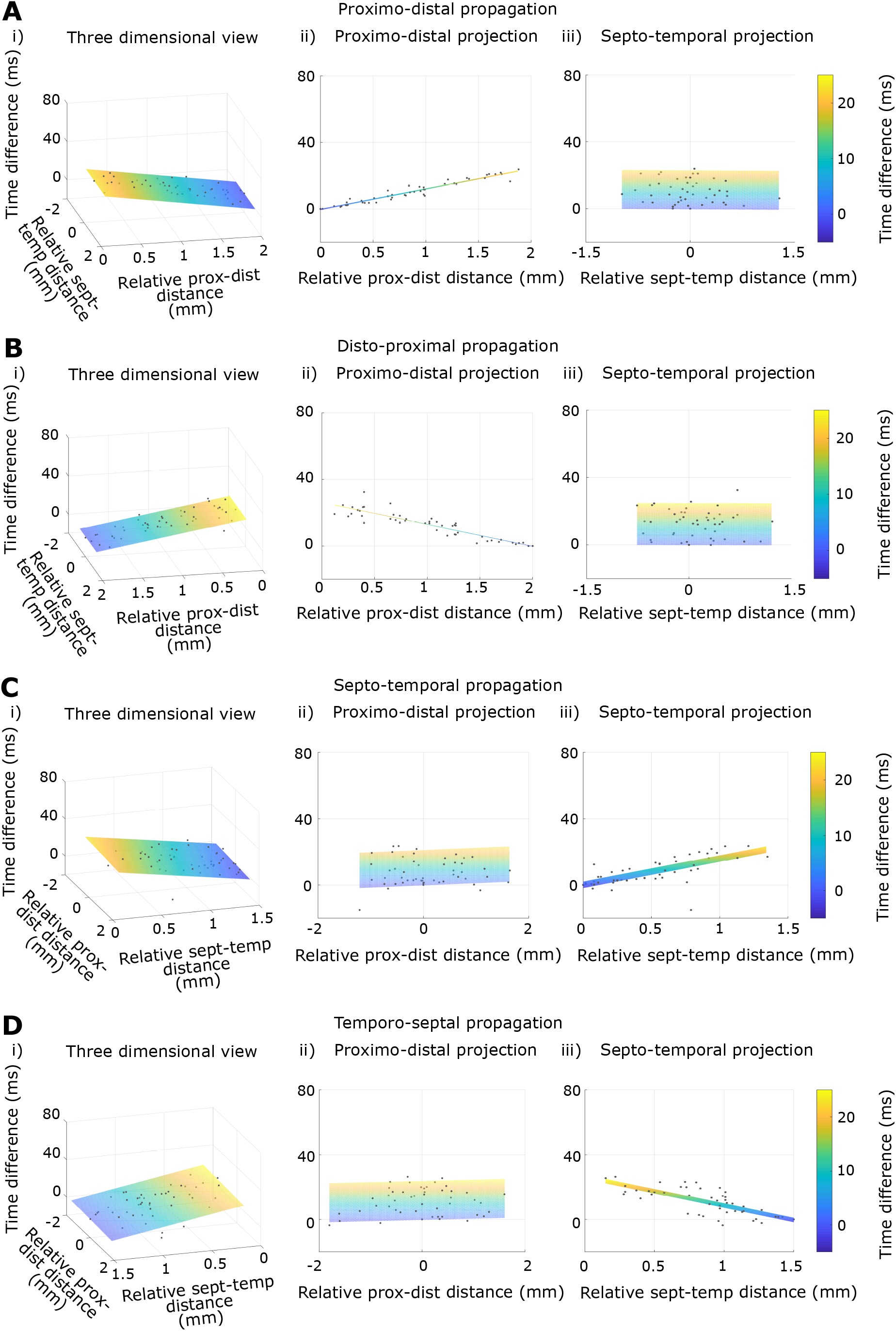
Ripple propagation speed for ≥ 2 SD. (A)-(D) Distributions of relative time difference vs. relative distances along the proximo-distal and septotemporal axes after classification of events into the four directions of propagation show clear narrow, linear trends along the expected direction of propagation. Column (i) shows a 3D (X-Y-Z) view, (ii) shows a proximo-distal projection (X-Z view), and (iii) shows a septo-temporal projection (Y-Z view) of column (i). Orientation of plots for propagation along the proximo-distal axis (Ai and Bi) are different from those along the septo-temporal axis (Ci and Di) for visualization purposes.

The propagation statistics remained similar after controlling for differences in the spatial spreads of tetrodes across the proximo-distal and septo-temporal axes (Supplementary Figure 12,13,14,15, and Supplementary Table 14).

## DISCUSSION

In this study, we show that lower amplitude ripples preferentially occurred towards proximal CA1 while higher amplitude ripples preferentially occurred towards distal CA1. CA3 activity, which is known to induce sharp-wave ripples in CA1 (Buzsàki et al., 1992; Csicsvari et al., 2000), could give rise to the observed differences in the amplitude of ripples. Distal CA1 receives stronger projections from proximal CA3 while proximal CA1 receives stronger projections from distal CA3 (Ishizuka et al., 1990). Thus, the higher firing rates of proximal CA3 (Csicsvari et al., 2000) may induce higher amplitude ripples in distal CA1. We further showed that the amplitude and the probability of co-occurrence of ripples decreased as a function of distance from the reference. This fall-off was more reliable along the proximo-distal axis than along a similar spatial extent of the septo-temporal axis. The relatively small decline in amplitude and probability of co-occurrence across our spatial range of the septo-temporal axis seem consistent with previous findings (Patel et al., 2013) within a similar spatial range. Additionally, in line with the Patel et al. (2013) study, we found that ripples propagate in CA1. They have shown that ripples propagate along a long range of the septo-temporal axis. We showed that ripples propagate along the proximo-distal and a comparable spatial extent of the septo-temporal axis. Furthermore, the ripples preferentially propagated along the proximo-distal axis (63% of the propagating ripple events propagated along the proximo-distal axis while 37% propagated along the septo-temporal axis).

The propagation speeds along both the proximo-distal (~0.08 mm/ms) and septo-temporal axes (~0.06 mm/ms) reported in this study are slower than the septo-temporal speed reported in the Patel et al. (2013) study (~0.35 mm/ms). These differences could arise because of possible methodological differences in the experiment and analyses between the two studies. Patel et al. (2013) used only the ripple events during sleep, while our dataset includes periods of awake immobility as well as sleep, as we did not explicitly record the behavioral state of the animal while it rested on the pedestal. Network components and dynamics involved in ripple generation and propagation may be different during different behavioral states (Buzsàki, 2015; Mizuseki and Miyawaki, 2017), giving rise to different speeds. A substantial majority of the events in our dataset propagate at speeds less than 0.2 mm/ms, and there does not appear to be a bimodal distribution with a secondary peak around 0.35 mm/ms (Supplementary Figure 16; fraction of propagating events with speeds less than 0.2 mm/ms = 0.876). Unless we have only a small minority of propagating ripple events recorded during sleep in our dataset, this lack of fast propagating events is indicative of ripple propagation speeds being indeed slower in our dataset irrespective of the behavioral state of the rat. We could not determine the criteria used for identifying propagating ripples from the description in Patel et al. (2013), but they plot pairwise temporal differences for speed estimation. In contrast, our study uses the ripple timing over the entire tetrode distribution to both identify propagating ripples as well as measure propagation speeds. Differences in criteria used to classify events as propagating could influence the estimated speed.

What mechanism might give rise to the observed preference in propagation along the proximo-distal axis versus the septo-temporal axis? The two most obvious candidates are 1. the CA3 projections inducing sharp-wave ripple complexes in CA1 (Buzsàki et al., 1992; Csicsvari et al., 2000) and 2. the interactions of CA1 pyramidal cells with basket cells and bistratified cells, the interneurons implicated in ripple generation/timing within CA1 (Ylinen et al., 1995; Csicsvari et al., 1999; Klausberger et al., 2004; Buzsàki, 2015). The topographical organization of CA3 to CA1 connectivity is complex, both in terms of anatomy and physiology (Ishizuka et al., 1990; Csicsvari et al., 2000). Population bursts in CA3 correlated with CA1 sharp-waves show propagation from distal to proximal CA3 (Csicsvari et al., 2000), predicting a directional preference in ripple propagation, not seen in our data. It is possible that the higher firing rates of proximal CA3 pyramidal cells they report compensate by ensuring their post-synaptic targets reach ripple generation threshold sooner in the sharp-wave in some of the events. Whether the CA3 population bursts propagate more along the proximo-distal axis than the septo-temporal axis is unknown. Even if that were the case, the CA3 to CA1 projection pattern is oblique, with proximal CA3 preferentially projecting more septally to distal CA1. Conversely, distal CA3 preferentially projects more temporally to proximal CA1 (Ishizuka et al., 1990). This CA3 to CA1 connectivity would predict an oblique spread of ripples within CA1 in response to CA3 activation propagating preferentially along the proximo-distal axis, if CA3 to CA1 connectivity alone were driving CA1 ripple propagation. Basket cell and bistratified cell axonal distributions (Sik et al., 1995, Klausberger et al., 2004) appear to be symmetric and show ~1 mm spread in both proximo-distal and septo-temporal directions. Hence, the interneuron connectivity could possibly explain the non-preferential propagation of the lower amplitude ripples. However, interneuron activity alone cannot account for the proximo-distal vs. septo-temporal difference in ripple propagation of higher amplitude ripples.

Even if the CA1 circuitry responsible for ripple generation is symmetric, asymmetric inputs can bias the starting point of propagating ripples to their target locations. Such asymmetry in inputs can lead to the observed anisotropy in preferred axes of ripple propagation. There are multiple potential candidates that may account for input asymmetry. The dentate gyrus (Sasaki et al., 2018; Sullivan et al., 2011), CA2 (Oliva et al., 2016a), and GABAergic projections from the medial septum (Dragoi et al., 1999; Viney et al., 2013) have been implicated in sharp-wave ripple initiation. Paucity of data about the projection patterns of these regions onto CA1 restricts our ability to understand their possible role in our observations.

LEC and MEC show differential projections to CA1 along the proximo-distal axis (Steward and Scoville, 1976; Naber et al., 2001). These differential projections could possibly explain the preferential propagation along the proximo-distal axis over the septo-temporal axis. There is some evidence for MEC activation during ripples. MEC shows high frequency events associated with ripples (Chrobak and Buzsàki, 1996). Buzsàki (2015), using data from Mizuseki et al. (2009), showed that superficial MEC activity peaks after the ripple peak, but the activity starts slowly ramping up some time before the peak (but see Chrobak and Buzsàki, 1994; O’Neill et al., 2017). Sullivan et al. (2011) showed an increase in superficial MEC firing correlated with increased probability of ripples at the DOWN to UP state transition. In mice, MEC LIII activity is essential for ripple bursts in CA1 during awake immobility (Yamamoto and Tonegawa, 2017). To our knowledge, LEC activity during CA1 ripples has not been studied. If MEC and LEC are independently active during a subset of ripples, they may bias the ripples to start earlier in proximal or distal CA1 respectively, leading to the preferential propagation along the proximo-distal axis observed in this study. Further studies are, however, required to ascertain the contribution of the various inputs described above to the propagation of CA1 ripples.

The biasing of a relatively large number of CA1 propagating ripples along the proximo-distal axis could imply that transfer of information to the cortical areas for consolidation happens in an orderly fashion, with some events beginning with spatial (where) information and ending with sensory (what/landmark related spatial) information, while others beginning with sensory information and ending with spatial information. This biased CA1 ripple propagation would, in turn, add a layer of complexity to the reciprocal communication between the hippocampus and the cortex (Peyrache et al., 2009; Rothschild et al., 2016).

In conclusion, our study shows a difference in ripple propagation along the proximo-distal and the septo-temporal axes, suggesting an anisotropy in the neural network along the two axes. Further studies are essential to discern the underlying mechanisms and functional significance of this observation.

## Acknowledgements

We thank Jeremy Johnson, Geeta Rao, Vyash Puliyadi, Amanda Smolinsky, Lou Blanpain, and Kimberley Christian for their support in data collection; Aditi Bishnoi, Indraja Jakhalekar and Rajat Saxena for their inputs on the manuscript. This work was supported by Wellcome Trust/DBT India Alliance Grant IA/S/13/2/501024. Data collection was supported by R01 NS039456 (J. Knierim, PI).

