## SupplementaryTablesandFigures for "Differential propagation of ripples along the proximo-distal and septo-temporal axes of dorsal CA1 of rats"

**SUPPLEMENTARY INFORMATION**

**Supplementary Methods**

Analysis controlling for the direction from the reference to the referred tetrode

Based on the direction from the reference to the referred tetrodes along the proximo-distal and septo-temporal axes, the data for co-occurrence and relative amplitude were each separated into four groups – tetrode pairs having more proximal and septal reference tetrodes than their corresponding referred tetrodes (PD ST), tetrode pairs having more distal and septal reference tetrodes than their corresponding referred tetrodes (DP ST), tetrode pairs having more proximal and temporal reference tetrodes than their corresponding referred tetrodes (PD TS), and tetrode pairs having more distal and temporal reference tetrodes than their corresponding referred tetrodes (DP TS). This was done to ensure that there was no influence of directionality.

Analysis using comparable spatial spreads

To ascertain that our results were not influenced by the differences in the spatial spread of the tetrodes across the proximo-distal and septo-temporal axes, the ripple co-occurrence, amplitude and propagation analyses were repeated by eliminating tetrodes to make the spatial spread along the two axes comparable. The elimination was done so as to maximize the spread along the septo-temporal axis and minimize the difference in the spatial spread along the two axes. At most three tetrodes were eliminated from each rat. Seven rats were utilized for the co-occurrence and amplitude analyses under this condition (for rat 302, no combination of tetrode elimination yielded a comparable spatial spread across the two axes even for the required minimum two tetrodes because of which it was eliminated from the analysis). For the propagation analysis, five rats were utilized as the remaining three (rat 302, rat 305, rat 392) did not satisfy the criterion of having a minimum of six tetrodes after eliminating tetrodes.

**Supplementary Tables**

| Amplitude group | Proximo-distal | | Septo-temporal | | p-value of comparison of slopes |
|---|---|---|---|---|---|
| | **partial slope** | **p-value** | **partial slope** | **p-value** | |
| 2-4 SD | -2.58 | 0.04* | -0.49 | 0.79 | 0.17 |
| 4-6 SD | -0.65 | 0.42 | -0.93 | 0.42 | 0.58 |
| ≥ 6 SD | 2.06 | 0.011* | -0.073 | 0.95 | 0.084 |

**Supplementary Table 1** Rate of ripple occurrence.

Change of rate of ripple occurrence as a function of relative position (count/min/mm) along the proximo-distal and septo-temporal axes for 2-4 SD, 4-6 SD, and ≥ 6 SD.

| Rat number | Amplitude group | Proximo-distal | | Septo-temporal | | p-value of comparison of slopes |
|---|---|---|---|---|---|---|
| | | partial slope | p-value | partial slope | p-value | |
| **302** | ≥ 2 SD | -9.2 | 0.03* | 1.97 | 0.77 | 0.22 |
| | 2-4 SD | -8.99 | 0.02* | 1.8 | 0.76 | 0.19 |
| | 4-6 SD | -3.4 | 0.02* | 0.76 | 0.74 | 0.20 |
| | ≥ 6 SD | 3.21 | 0.005* | -0.60 | 0.65 | 0.096 |
| **305** | ≥ 2 SD | -6.1 | 0.37 | 4.8 | 0.65 | 0.46 |
| | 2-4 SD | -7.03 | 0.3 | 4.3 | 0.63 | 0.39 |
| | 4-6 SD | -2.66 | 0.36 | 2.04 | 0.64 | 0.45 |
| | ≥ 6 SD | 3.64 | 0.18 | -1.54 | 0.59 | 0.27 |
| **391** | ≥ 2 SD | -3.17 | 0.21 | -1.71 | 0.62 | 0.37 |
| | 2-4 SD | -4.4 | 0.03* | -0.43 | 0.84 | 0.11 |
| | 4-6 SD | -1.22 | 0.19 | -0.87 | 0.49 | 0.41 |
| | ≥ 6 SD | 2.48 | 0.0014* | -0.41 | 0.43 | 0.13 |
| **392** | ≥ 2 SD | -2.6 | 0.4 | -4.23 | 0.3 | 0.66 |
| | 2-4 SD | -2.97 | 0.21 | -3.84 | 0.22 | 0.62 |
| | 4-6 SD | -1.14 | 0.33 | -2.33 | 0.16 | 0.78 |
| | ≥ 6 SD | 1.44 | 0.15 | 1.93 | 0.14 | 0.67 |
| **416** | ≥ 2 SD | -1.72 | 0.45 | -9.08 | 0.09 | 0.88 |
| | 2-4 SD | -4.08 | 0.02* | -3.42 | 0.23 | 0.42 |
| | 4-6 SD | -1.004 | 0.17 | -3.11 | 0.06 | 0.87 |
| | ≥ 6 SD | 3.35 | 0.0069* | -2.56 | 0.16 | 0.35 |
| **417** | ≥ 2 SD | -2.4 | 0.4 | 4.3 | 0.2 | 0.67 |
| | 2-4 SD | -3.03 | 0.09 | 5.27 | 0.02* | 0.97 |
| | 4-6 SD | -2.32 | 0.08 | 1.54 | 0.24 | 0.33 |
| | ≥ 6 SD | 3.00 | $6 \times 10^{-4}$* | -2.53 | 0.002* | 0.23 |
| **432** | ≥ 2 SD | 0.43 | 0.66 | 2.74 | 0.1 | 0.89 |
| | 2-4 SD | 0.07 | 0.89 | 0.22 | 0.03* | 0.97 |
| | 4-6 SD | -0.05 | 0.85 | 1.1 | 0.02* | 0.98 |
| | ≥ 6 SD | 0.42 | 0.52 | -0.67 | 0.49 | 0.59 |
| **441** | ≥ 2 SD | 2.2 | 0.6 | 5.5 | 0.4 | 0.71 |
| | 2-4 SD | -0.3 | 0.92 | 1.83 | 0.72 | 0.63 |
| | 4-6 SD | 0.68 | 0.56 | 1.86 | 0.34 | 0.75 |
| | ≥ 6 SD | 1.87 | 0.015* | 1.81 | 0.092 | 0.47 |

**Supplementary Table 2** Rat-wise rate of ripple occurrence.

Change of rate of ripple occurrence as a function of relative position (count/min/mm) along the proximo-distal and septo-temporal axes for ≥ 2 SD, 2-4 SD, 4-6 SD, and ≥ 6 SD for each rat separately. Consistent with the average data, a majority of the rats showed opposing trends between 2-4 SD and ≥ 6 SD, though they were not always significant. Each rat has a corresponding colored dot in Figure 1B: black – rat 302, red – rat 305, dark blue – rat 391, sea green – rat 392, light blue – rat 416, lime green – rat 417, purple – rat 432, pink – rat 441.

| Amplitude group | Proximo-distal | | Septo-temporal | | p-value of comparison of slopes |
|---|---|---|---|---|---|
| | partial slope | p-value | partial slope | p-value | |
| ≥ 2 SD | 0.09 | 0.97 | -0.21 | 0.95 | 0.51 |
| 2-4 SD | -0.19 | 0.91 | 0.34 | 0.86 | 0.52 |
| 4-6 SD | -0.25 | 0.83 | -0.49 | 0.71 | 0.55 |
| ≥ 6 SD | 0.57 | 0.59 | -0.039 | 0.97 | 0.37 |

**Supplementary Table 3** Rate of ripple occurrence for comparable spatial spreads.

Change of rate of ripple occurrence as a function of relative position (count/min/mm) along the proximo-distal and septo-temporal axes for ≥ 2 SD, 2-4 SD, 4-6 SD, and ≥ 6 SD after controlling for differences in spatial spreads of tetrodes along the proximo-distal and septo-temporal axes.

| Amplitude group | Proximo-distal | | Septo-temporal | | p-value of comparison of slopes |
|---|---|---|---|---|---|
| | partial slope | p-value | partial slope | p-value | |
| 2-4 SD | -0.069 | $4 \times 10^{-10}$* | -0.022 | 0.12 | 0.003* |
| 4-6 SD | -0.088 | $2 \times 10^{-20}$* | -0.032 | 0.006* | $3 \times 10^{-5}$* |
| ≥ 6 SD | -0.039 | $1 \times 10^{-24}$* | -0.017 | $3 \times 10^{-4}$* | $2 \times 10^{-5}$* |

**Supplementary Table 4** Ripple co-occurrence.

Change of fraction of co-occurring ripples as a function of relative distance between tetrode pairs (fraction of co-occurring ripples/mm) along the proximo-distal and septo-temporal axes for 2-4 SD, 4-6 SD, and ≥ 6 SD.

| Rat number | Amplitude group | Proximo-distal | | Septo-temporal | | p-value of comparison of slopes |
|---|---|---|---|---|---|---|
| | | partial slope | p-value | partial slope | p-value | |
| **302** | ≥ 2 SD | -0.093 | 0.002* | 0.089 | 0.16 | 0.48 |
| | 2-4 SD | -0.11 | $2 \times 10^{-4}$* | 0.075 | 0.18 | 0.28 |
| | 4-6 SD | -0.12 | $2 \times 10^{-5}$* | 0.087 | 0.08 | 0.27 |
| | ≥ 6 SD | -0.04 | $6 \times 10^{-6}$* | 0.02 | 0.16 | 0.12 |
| **305** | ≥ 2 SD | -0.17 | 0.14 | -0.046 | 0.71 | 0.14 |
| | 2-4 SD | -0.16 | 0.12 | -0.046 | 0.67 | 0.14 |
| | 4-6 SD | -0.18 | 0.13 | -0.052 | 0.69 | 0.14 |
| | ≥ 6 SD | -0.089 | 0.08 | -0.013 | 0.79 | 0.075 |
| **391** | ≥ 2 SD | -0.08 | $1 \times 10^{-5}$* | 0.027 | 0.18 | 0.0096* |
| | 2-4 SD | -0.13 | $2 \times 10^{-5}$* | 0.025 | 0.44 | 0.003* |
| | 4-6 SD | -0.11 | $7 \times 10^{-6}$* | 0.037 | 0.15 | 0.007* |
| | ≥ 6 SD | -0.024 | $6 \times 10^{-5}$* | 0.0077 | 0.25 | 0.015* |
| **392** | ≥ 2 SD | -0.03 | 0.05 | -0.0011 | 0.96 | 0.13 |
| | 2-4 SD | -0.029 | 0.063 | -0.010 | 0.66 | 0.23 |
| | 4-6 SD | -0.039 | 0.014* | -0.029 | 0.22 | 0.35 |
| | ≥ 6 SD | -0.025 | $2 \times 10^{-5}$* | -0.0044 | 0.54 | 0.0065* |
| **416** | ≥ 2 SD | -0.05 | $2 \times 10^{-5}$* | -0.084 | $8 \times 10^{-5}$* | 0.96 |
| | 2-4 SD | -0.045 | 0.0014* | -0.11 | 0.0001* | 0.99 |
| | 4-6 SD | -0.079 | $3 \times 10^{-9}$* | -0.10 | $3 \times 10^{-6}$* | 0.9 |
| | ≥ 6 SD | -0.033 | $3 \times 10^{-7}$* | -0.04 | $5 \times 10^{-5}$* | 0.9 |
| **417** | ≥ 2 SD | -0.07 | $2 \times 10^{-6}$* | -0.05 | $2 \times 10^{-4}$* | 0.07 |
| | 2-4 SD | -0.046 | $4 \times 10^{-7}$* | -0.052 | $3 \times 10^{-8}$* | 0.74 |
| | 4-6 SD | -0.093 | $1 \times 10^{-6}$* | -0.049 | 0.0016* | 0.014* |
| | ≥ 6 SD | -0.064 | $7 \times 10^{-9}$* | -0.023 | 0.003* | $1 \times 10^{-4}$* |
| **432** | ≥ 2 SD | -0.06 | $2 \times 10^{-7}$* | -0.023 | 0.08 | 0.007* |
| | 2-4 SD | -0.073 | $2 \times 10^{-4}$* | -0.034 | 0.18 | 0.072 |
| | 4-6 SD | -0.084 | $2 \times 10^{-10}$* | -0.04 | 0.0082* | 0.0013* |
| | ≥ 6 SD | -0.025 | $1 \times 10^{-10}$* | -0.0098 | 0.018* | $9 \times 10^{-10}$* |
| **441** | ≥ 2 SD | -0.089 | $3 \times 10^{-7}$* | -0.007 | 0.72 | $7 \times 10^{-4}$* |
| | 2-4 SD | -0.071 | $1 \times 10^{-5}$* | -0.022 | 0.27 | 0.021* |
| | 4-6 SD | -0.10 | $7 \times 10^{-8}$* | -0.022 | 0.31 | 0.0014* |
| | ≥ 6 SD | -0.049 | $9 \times 10^{-10}$* | -0.01 | 0.24 | $2 \times 10^{-4}$* |

**Supplementary Table 5** Rat-wise ripple co-occurrence.

Change of fraction of co-occurring ripples as a function of relative distance between tetrode pairs (fraction of co-occurring ripples/mm) along the proximo-distal and septo-temporal axes for ≥ 2 SD, 2-4 SD, 4-6 SD, and ≥ 6 SD for each rat separately. Consistent with the average data, 7/8 rats showed a significant decrease along the proximo-distal axis in two or more amplitude groups, while the decrease along the septo-temporal axis was not so reliable across rats.

| Amplitude group | Direction from reference to referred | Proximo-distal | | Septo-temporal | | p-value of comparison of slopes |
|---|---|---|---|---|---|---|
| | | partial slope | p-value | partial slope | p-value | |
| ≥ 2 SD | PD ST | -0.093 | $2 \times 10^{-6}$* | -0.039 | 0.14 | 0.033* |
| | DP ST | -0.076 | $7 \times 10^{-7}$* | 0.0064 | 0.82 | 0.0016* |
| | PD TS | -0.078 | $2 \times 10^{-5}$* | 0.0083 | 0.73 | 0.0093* |
| | DP TS | -0.048 | 0.0016* | -0.044 | 0.038* | 0.43 |
| 2-4 SD | PD ST | -0.074 | $2 \times 10^{-4}$* | -0.023 | 0.39 | 0.052 |
| | DP ST | -0.089 | $1 \times 10^{-5}$* | -0.013 | 0.62 | 0.0093* |
| | PD TS | -0.079 | $2 \times 10^{-4}$* | -0.01 | 0.72 | 0.021* |
| | DP TS | -0.029 | 0.13 | -0.019 | 0.49 | 0.36 |
| 4-6 SD | PD ST | -0.086 | $4 \times 10^{-8}$* | -0.051 | 0.013* | 0.063 |
| | DP ST | -0.11 | $6 \times 10^{-10}$* | -0.0017 | 0.93 | $4 \times 10^{-5}$* |
| | PD TS | -0.092 | $3 \times 10^{-8}$* | -0.025 | 0.24 | 0.0044* |
| | DP TS | -0.074 | $6 \times 10^{-6}$* | -0.038 | 0.08 | 0.074 |
| ≥ 6 SD | PD ST | -0.034 | $3 \times 10^{-10}$* | -0.025 | $5 \times 10^{-4}$* | 0.11 |
| | DP ST | -0.049 | $8 \times 10^{-13}$* | -0.0044 | 0.59 | $8 \times 10^{-6}$* |
| | PD TS | -0.033 | $2 \times 10^{-9}$* | -0.012 | 0.094 | 0.005* |
| | DP TS | -0.045 | $2 \times 10^{-11}$* | -0.023 | 0.0069* | 0.0095* |

**Supplementary Table 6** Ripple co-occurrence accounting for the direction from the reference to the referred tetrode.

Change of fraction of co-occurring ripples as a function of relative distance between tetrode pairs (fraction of co-occurring ripples/mm) along the proximo-distal and septo-temporal axes for ≥ 2 SD, 2-4 SD, 4-6 SD, and ≥ 6 SD after controlling for the direction from the reference to the referred tetrode along the two axes (as described in Supplementary Methods). PD denotes the reference tetrode being more proximal to the referred tetrode, DP denotes the reference tetrode being more distal to the referred tetrode, ST denotes the reference tetrode being more septal to the referred tetrode, while TS denotes the reference tetrode being more temporal to the referred tetrode.

| **Amplitude group** | **Proximo-distal** | | **Septo-temporal** | | **p-value of comparison of slopes** |
|---|---|---|---|---|---|
| | **partial slope** | **p-value** | **partial slope** | **p-value** | |
| ≥ 2 SD | -0.058 | 0.0012* | -0.025 | 0.14 | 0.088 |
| 2-4 SD | -0.077 | $8 \times 10^{-4}$* | -0.026 | 0.22 | 0.052 |
| 4-6 SD | -0.081 | $3 \times 10^{-6}$* | -0.031 | 0.047* | 0.015* |
| ≥ 6 SD | -0.028 | $8 \times 10^{-7}$* | -0.015 | 0.003* | 0.046* |

**Supplementary Table 7** Ripple co-occurrence for comparable spatial spreads.

Change of fraction of co-occurring ripples as a function of relative distance between tetrode pairs (fraction of co-occurring ripples/mm) along the proximo-distal and septo-temporal axes for ≥ 2 SD, 2-4 SD, 4-6 SD, and ≥ 6 SD after controlling for differences in spatial spreads of tetrodes along the proximo-distal and septo-temporal axes.

| Amplitude group | Proximo-distal | | Septo-temporal | | p-value of comparison of slopes |
|---|---|---|---|---|---|
| | partial slope | p-value | partial slope | p-value | |
| 2-4 SD | $6 \times 10^{-5}$ | 0.99 | 0.0099 | 0.32 | 0.8 |
| 4-6 SD | -0.0048 | 0.44 | -0.0006 | 0.94 | 0.34 |
| ≥ 6 SD | -0.057 | $5 \times 10^{-21}$* | -0.031 | $7 \times 10^{-5}$* | 0.0012* |

**Supplementary Table 8** Relative ripple amplitude.

Change of relative ripple amplitude as a function of relative distance between tetrode pairs (relative amplitude/mm) along the proximo-distal and septo-temporal axes for 2-4 SD, 4-6 SD, and ≥ 6 SD.

| Rat number | Amplitude group | Proximo-distal | | Septo-temporal | | p-value of comparison of slopes |
|---|---|---|---|---|---|---|
| | | partial slope | p-value | partial slope | p-value | |
| **302** | ≥ 2 SD | -0.04 | 0.0011* | 0.029 | 0.25 | 0.35 |
| | 2-4 SD | -0.0068 | 0.46 | 0.011 | 0.65 | 0.55 |
| | 4-6 SD | -0.007 | 0.49 | 0.019 | 0.46 | 0.65 |
| | ≥ 6 SD | -0.056 | $1 \times 10^{-4}$* | 0.049 | 0.08 | 0.42 |
| **305** | ≥ 2 SD | -0.055 | 0.10 | -0.016 | 0.65 | 0.12 |
| | 2-4 SD | -0.046 | 0.075 | -0.015 | 0.56 | 0.11 |
| | 4-6 SD | -0.037 | 0.34 | -0.017 | 0.72 | 0.32 |
| | ≥ 6 SD | -0.11 | 0.062 | -0.027 | 0.62 | 0.077 |
| **391** | ≥ 2 SD | -0.012 | 0.02* | 0.009 | 0.17 | 0.36 |
| | 2-4 SD | 0.029 | 0.069 | 0.014 | 0.51 | 0.27 |
| | 4-6 SD | 0.03 | 0.013* | 0.004 | 0.78 | 0.07 |
| | ≥ 6 SD | -0.05 | $2 \times 10^{-6}$* | 0.0077 | 0.5 | $5 \times 10^{-4}$* |
| **392** | ≥ 2 SD | -0.0034 | 0.56 | 0.0066 | 0.46 | 0.63 |
| | 2-4 SD | 0.015 | 0.10 | -0.029 | 0.054 | 0.81 |
| | 4-6 SD | 0.033 | $2 \times 10^{-4}$* | 0.015 | 0.18 | 0.077 |
| | ≥ 6 SD | -0.038 | $5 \times 10^{-5}$* | -0.006 | 0.58 | 0.01* |
| **416** | ≥ 2 SD | -0.015 | 0.15 | -0.05 | 0.009* | 0.97 |
| | 2-4 SD | 0.025 | 0.016* | -0.029 | 0.14 | 0.57 |
| | 4-6 SD | -0.002 | 0.77 | -0.002 | 0.87 | 0.5 |
| | ≥ 6 SD | -0.05 | $3 \times 10^{-8}$* | -0.056 | $6 \times 10^{-5}$* | 0.73 |
| **417** | ≥ 2 SD | -0.037 | $1 \times 10^{-6}$* | -0.018 | 0.003* | 0.011* |
| | 2-4 SD | 0.001 | 0.85 | -0.017 | 0.0037* | 0.98 |
| | 4-6 SD | -0.043 | 0.003* | -0.0032 | 0.8 | 0.013* |
| | ≥ 6 SD | -0.087 | $1 \times 10^{-8}$* | -0.038 | 0.001* | $6 \times 10^{-4}$* |
| **432** | ≥ 2 SD | -0.0016 | 0.90 | 0.037 | 0.063 | 0.96 |
| | 2-4 SD | 0.016 | 0.009* | 0.02 | 0.023* | 0.68 |
| | 4-6 SD | 0.0094 | 0.011* | 0.015 | 0.0054* | 0.86 |
| | ≥ 6 SD | -0.044 | $4 \times 10^{-11}$* | -0.016 | 0.024* | $1 \times 10^{-4}$* |
| **441** | ≥ 2 SD | -0.038 | $2 \times 10^{-5}$* | -0.0015 | 0.89 | 0.0038* |
| | 2-4 SD | -0.015 | 0.23 | -0.006 | 0.73 | 0.33 |
| | 4-6 SD | -0.0066 | 0.56 | 0.0022 | 0.89 | 0.41 |
| | ≥ 6 SD | -0.07 | $7 \times 10^{-11}$* | -0.0024 | 0.82 | $3 \times 10^{-6}$* |

**Supplementary Table 9** Rat-wise relative ripple amplitude.

Change of relative ripple amplitude as a function of relative distance between tetrode pairs (relative amplitude/mm) along the proximo-distal and septo-temporal axes for ≥ 2 SD, 2-4 SD, 4-6 SD, and ≥ 6 SD for each rat separately. Consistent with the average data, for ≥ 2 SD, 4 rats showed a significant decrease along the proximo-distal axis and none of the rats showed the opposite trend. Relative amplitude for 2-4 SD and 4-6 SD did not show any gradients along the proximo-distal or septo-temporal axes in the average data, however, the rat-wise data showed a variation of positive and negative gradients among the different rats. Consistent with the average data, for ≥ 6 SD the decrease along the proximo-distal axis was significant for 7/8 rats, while the decrease along the septo-temporal axis was not as reliable across rats.

| **Amplitude group** | **Direction from reference to referred** | **Proximo-distal** | | **Septo-temporal** | | **p-value of comparison of slopes** |
|---|---|---|---|---|---|---|
| | | **partial slope** | **p-value** | **partial slope** | **p-value** | |
| ≥ 2 SD | PD ST | -0.043 | 0.10 | 0.014 | 0.70 | 0.23 |
| | DP ST | -0.0043 | 0.84 | -0.035 | 0.25 | 0.79 |
| | PD TS | -0.024 | 0.25 | 0.037 | 0.19 | 0.64 |
| | DP TS | -0.024 | 0.21 | -0.027 | 0.31 | 0.54 |
| 2-4 SD | PD ST | -0.04 | 0.09 | 0.005 | 0.87 | 0.16 |
| | DP ST | 0.04 | 0.12 | -0.023 | 0.55 | 0.33 |
| | PD TS | 0.014 | 0.62 | 0.085 | 0.028* | 0.93 |
| | DP TS | 0.011 | 0.71 | -0.004 | 0.91 | 0.44 |
| 4-6 SD | PD ST | -0.019 | 0.25 | -0.017 | 0.46 | 0.46 |
| | DP ST | 0.025 | 0.13 | -0.032 | 0.16 | 0.6 |
| | PD TS | 0.0045 | 0.84 | 0.044 | 0.16 | 0.84 |
| | DP TS | -0.019 | 0.28 | -0.0053 | 0.83 | 0.3 |
| ≥ 6 SD | PD ST | -0.057 | $3 \times 10^{-7}$* | -0.031 | 0.038* | 0.05 |
| | DP ST | -0.066 | $6 \times 10^{-6}$* | -0.036 | 0.06 | 0.10 |
| | PD TS | -0.053 | $4 \times 10^{-4}$* | -0.016 | 0.44 | 0.066 |
| | DP TS | -0.069 | $2 \times 10^{-7}$* | -0.034 | 0.048* | 0.037* |

**Supplementary Table 10** Relative ripple amplitude accounting for the direction from the reference to the referred tetrode.

Change of relative ripple amplitude as a function of relative distance between tetrode pairs (fraction of co-occurring ripples/mm) along the proximo-distal and septo-temporal axes for ≥ 2 SD, 2-4 SD, 4-6 SD, and ≥ 6 SD after controlling for the direction from the reference to the referred tetrode along the two axes (as described in Supplementary Methods). PD denotes the reference tetrode being more proximal to the referred tetrode, DP denotes the reference tetrode being more distal to the referred tetrode, ST denotes the reference tetrode being more septal to the referred tetrode, while TS denotes the reference tetrode being more temporal to the referred tetrode.

| **Amplitude group** | **Proximo-distal** | | **Septo-temporal** | | **p-value of comparison of slopes** |
|---|---|---|---|---|---|
| | **partial slope** | **p-value** | **partial slope** | **p-value** | |
| ≥ 2 SD | 0.002 | 0.87 | -0.005 | 0.61 | 0.59 |
| 2-4 SD | 0.021 | 0.18 | 0.0013 | 0.93 | 0.18 |
| 4-6 SD | 0.031 | 0.021* | -0.004 | 0.76 | 0.072 |
| ≥ 6 SD | -0.035 | 0.0033* | -0.029 | 0.011* | 0.35 |

**Supplementary Table 11** Relative ripple amplitude for comparable spatial spreads.

Change of relative ripple amplitude as a function of relative distance between tetrode pairs (relative amplitude/mm) along the proximo-distal and septo-temporal axes for ≥ 2 SD, 2-4 SD, 4-6 SD, and ≥ 6 SD after controlling for differences in spatial spreads of tetrodes along the proximo-distal and septo-temporal axes.

| Amplitude group | Direction of propagation | Number of ripples | Proximo-distal | | Septo-temporal | | p-value of comparison of slopes | Resultant speed |
|---|---|---|---|---|---|---|---|---|
| | | | partial slope | p-value | partial slope | p-value | | |
| 2-4 SD | Prox-dist | 139 (30%) | 32.4 | $1 \times 10^{-20}$* | 0.51 | 0.86 | $4 \times 10^{-13}$* | 0.031 |
| | Dist-prox | 117 (25%) | -25.3 | $1 \times 10^{-16}$* | 0.84 | 0.77 | $8 \times 10^{-9}$* | 0.039 |
| | Sept-temp | 118(25%) | -0.65 | 0.64 | 33.9 | $8 \times 10^{-19}$* | 0* | 0.029 |
| | Temp-sept | 94 (20%) | 4.25 | 0.081 | -34.7 | $1 \times 10^{-9}$* | $1 \times 10^{-7}$* | 0.029 |
| 4-6 SD | Prox-dist | 300 (37%) | 16.9 | $1 \times 10^{-20}$* | -1.59 | 0.29 | $5 \times 10^{-12}$* | 0.059 |
| | Dist-prox | 173 (22%) | -25.2 | $6 \times 10^{-23}$* | -1.89 | 0.35 | $9 \times 10^{-13}$* | 0.039 |
| | Sept-temp | 150 (19%) | -2.13 | 0.16 | 26.6 | $2 \times 10^{-13}$* | $1 \times 10^{-11}$* | 0.037 |
| | Temp-sept | 175 (22%) | 2.6 | 0.11 | -20.7 | $3 \times 10^{-8}$* | $2 \times 10^{-6}$* | 0.048 |
| ≥ 6 SD | Prox-dist | 1286 (33%) | 10.02 | $1 \times 10^{-34}$* | 0.086 | 0.84 | 0* | 0.099 |
| | Dist-prox | 1269 (32%) | -11.3 | $1 \times 10^{-25}$* | 0.18 | 0.82 | $1 \times 10^{-15}$* | 0.089 |
| | Sept-temp | 704 (18%) | 1.45 | 0.19 | 12.8 | $3 \times 10^{-8}$* | $1 \times 10^{-6}$* | 0.078 |
| | Temp-sept | 674 (17%) | 0.52 | 0.55 | -15.4 | $5 \times 10^{-12}$* | $1 \times 10^{-10}$* | 0.065 |

**Supplementary Table 12** Ripple propagation.

Statistics for ripple propagation along the proximo-distal and septo-temporal axes for 2-4 SD, 4-6 SD, and ≥ 6 SD. Partial slopes in this case are the difference in time of occurrence of ripples as a function of distance along the proximo-distal and septo-temporal axes. The unit of the partial slopes is ms/mm while that of resultant speeds is mm/ms.

2-4 SD: number of putative events = 2579; number of propagating events = 468; percentage of propagating events = 18%; ratio for direction = 1.21; ratio for slopes = 0.833; test of proportions comparing the observed proportion of events with the 50% propagation along the proximo-distal axis expected by chance, Z = 2, p = 0.02*.

4-6 SD: number of putative events = 4484; number of propagating events = 798; percentage of propagating events = 18%; ratio for direction = 1.46; ratio for slopes = 0.418; test of proportions comparing the observed proportion of events with the 50% propagation along the proximo-distal axis expected by chance, Z = 5, p = $1 \times 10^{-7}$*.

≥ 6 SD: number of putative events = 18219; number of propagating events = 3933; percentage of propagating events = 21%; ratio for direction = 1.84; ratio for slopes = 0.328; test of proportions comparing the observed proportion of events with the 50% propagation along the proximo-distal axis expected by chance, $Z = 19$, $p = 0$*.

| Amplitude group | Direction of propagation | Number of ripples | Proximo-distal | | Septo-temporal | | p-value of comparison of slopes | Resultant speed |
|---|---|---|---|---|---|---|---|---|
| | | | partial slope | p-value | partial slope | p-value | | |
| ≥ 2 SD | Prox-dist | 83 (41%) | 11.5 | 0.0071* | -3.90 | 0.46 | 0.15 | 0.082 |
| | Dist-prox | 75 (37%) | -17.2 | 0.013* | 4.63 | 0.63 | 0.17 | 0.056 |
| | Sept-temp | 21 (11%) | -4.42 | 0.019* | 12.9 | 0.015* | 0.042* | 0.073 |
| | Temp-sept | 21 (11%) | 10.9 | 0.0022* | -29.2 | 0.0022* | 0.0092* | 0.032 |
| 2-4 SD | Prox-dist | 8 (35%) | 28.3 | 0.022* | -8.30 | 0.66 | 0.22 | 0.034 |
| | Dist-prox | 8 (35%) | -25.1 | 0.034* | 7.31 | 0.71 | 0.25 | 0.038 |
| | Sept-temp | 4 (17%) | 0.44 | 0.36 | 7.64 | 0.0062* | 0.0078* | 0.13 |
| | Temp-sept | 3 (13%) | 14.8 | 0.001* | -35.7 | 0.0014* | 0.0068* | 0.026 |
| 4-6 SD | Prox-dist | 13 (43%) | 14.5 | 0.0067* | -8.10 | 0.25 | 0.23 | 0.060 |
| | Dist-prox | 10 (33%) | -20.2 | 0.011* | 1.76 | 0.87 | 0.12 | 0.049 |
| | Sept-temp | 4 (13%) | -15.5 | 0.026* | 30.9 | 0.054 | 0.16 | 0.029 |
| | Temp-sept | 3 (10%) | 19.5 | 0.0042* | -34.01 | 0.014* | 0.095 | 0.025 |
| ≥ 6 SD | Prox-dist | 62 (42%) | 9.13 | 0.0047* | -2.82 | 0.44 | 0.11 | 0.10 |
| | Dist-prox | 57 (39%) | -15.3 | 0.011* | 4.74 | 0.55 | 0.17 | 0.062 |
| | Sept-temp | 13 (9%) | -2.53 | 0.01* | 9.01 | 0.0044* | 0.012* | 0.11 |
| | Temp-sept | 15 (10%) | 8.59 | 0.0048* | -22.3 | 0.0052* | 0.021* | 0.042 |

**Supplementary Table 13a** Ripple propagation for **rat 302.**

≥ 2 SD: number of putative events = 1365; number of propagating events = 200; percentage of propagating events = 15%; ratio for direction = 3.67; ratio for slopes = 0.115; test of proportions comparing the observed proportion of events with the 50% propagation along the proximo-distal axis expected by chance, Z = 8, p = 4 × $10^{-15}$*.

2-4 SD: number of putative events = 188; number of propagating events = 23; percentage of propagating events = 12%; ratio for direction = 2.28; ratio for slopes = 0; test of proportions comparing the observed proportion of events with the 50% propagation along the proximo-distal axis expected by chance, Z = 1.87, p = 0.03*.

4-6 SD: number of putative events = 297; number of propagating events = 30; percentage of propagating events = 10%; ratio for direction = 3.29; ratio for slopes = 0; test of proportions comparing the observed proportion of events with the 50% propagation along the proximo-distal axis expected by chance, Z = 3, p = 0.0017*.

≥ 6 SD: number of putative events = 880; number of propagating events = 147; percentage of propagating events = 17%; ratio for direction = 4.25; ratio for slopes = 0.130; test of proportions comparing the observed proportion of events with the 50% propagation along the proximo-distal axis expected by chance, Z = 7.5, p = $3 \times 10^{-13}$*.

| Amplitude group | Direction of propagation | Number of ripples | Proximo-distal | | Septo-temporal | | p-value of comparison of slopes | Resultant speed |
|---|---|---|---|---|---|---|---|---|
| | | | partial slope | p-value | partial slope | p-value | | |
| ≥ 2 SD | Prox-dist | 440 (40%) | 13.8 | $7 \times 10^{-5}$* | -1.57 | 0.26 | $7 \times 10^{-4}$* | 0.072 |
| | Dist-prox | 421 (39%) | -16.1 | $8 \times 10^{-5}$* | 0.44 | 0.78 | $6 \times 10^{-4}$* | 0.062 |
| | Sept-temp | 140 (13%) | 0.58 | 0.67 | 14.2 | 0.0018* | 0.0026* | 0.071 |
| | Temp-sept | 85 (8%) | 6.45 | 0.052 | -23.9 | 0.0025* | 0.0091* | 0.040 |
| 2-4 SD | Prox-dist | 17 (31%) | 48.3 | 0.0025* | -16.9 | 0.18 | 0.041* | 0.019 |
| | Dist-prox | 17 (31%) | -42.5 | $3 \times 10^{-5}$* | 2.34 | 0.49 | $3 \times 10^{-4}$* | 0.023 |
| | Sept-temp | 13 (24%) | 2.11 | 0.72 | 39.7 | 0.0079* | 0.011* | 0.025 |
| | Temp-sept | 7 (13%) | 22.05 | 0.077 | -79.8 | 0.0046* | 0.016* | 0.012 |
| 4-6 SD | Prox-dist | 73 (51%) | 15.4 | $2 \times 10^{-4}$* | -2.26 | 0.25 | 0.0018* | 0.064 |
| | Dist-prox | 35 (24%) | -27.7 | $3 \times 10^{-4}$* | -1.69 | 0.65 | 0.0021* | 0.036 |
| | Sept-temp | 24 (17%) | 1.85 | 0.034* | 11.9 | $2 \times 10^{-4}$* | $4 \times 10^{-4}$* | 0.082 |
| | Temp-sept | 12 (8%) | 18.8 | 0.0047* | -29.5 | 0.0039* | 0.085 | 0.029 |
| ≥ 6 SD | Prox-dist | 350 (39%) | 12.03 | $4 \times 10^{-5}$* | -0.43 | 0.66 | $3 \times 10^{-4}$* | 0.083 |
| | Dist-prox | 369 (42%) | -13.8 | $8 \times 10^{-5}$* | 0.64 | 0.64 | $6 \times 10^{-8}$* | 0.072 |
| | Sept-temp | 103 (12%) | -1.22 | 0.32 | 11.1 | 0.0024* | 0.0045* | 0.089 |
| | Temp-sept | 66 (7%) | 1.69 | 0.42 | -18.2 | 0.0029* | 0.0051* | 0.055 |

**Supplementary Table 13b** Ripple propagation for **rat 391.**

$\geq$ 2 SD: number of putative events = 6040; number of propagating events = 1086; percentage of propagating events = 18%; ratio for direction = 3.87; ratio for slopes = 0.108; test of proportions comparing the observed proportion of events with the 50% propagation along the proximo-distal axis expected by chance, $Z = 19$, $p = 0$*.

2-4 SD: number of putative events = 457; number of propagating events = 54; percentage of propagating events = 12%; ratio for direction = 1.70; ratio for slopes = 0; test of proportions comparing the observed proportion of events with the 50% propagation along the proximo-distal axis expected by chance, $Z = 1.9$, $p = 0.029$*.

4-6 SD: number of putative events = 976; number of propagating events = 144; percentage of propagating events = 15%; ratio for direction = 3; ratio for slopes = 0.083; test of proportions comparing the observed proportion of events with the 50% propagation along the proximo-distal axis expected by chance, $Z = 6$, $p = 2 \times 10^{-9}$*.

$\geq$ 6 SD: number of putative events = 4607; number of propagating events = 888; percentage of propagating events = 19%; ratio for direction = 4.25; ratio for slopes = 0.116; test of proportions comparing the observed proportion of events with the 50% propagation along the proximo-distal axis expected by chance, $Z = 18$, $p = 0$*.

| Amplitude group | Direction of propagation | Number of ripples | Proximo-distal | | Septo-temporal | | p-value of comparison of slopes | Resultant speed |
|---|---|---|---|---|---|---|---|---|
| | | | partial slope | p-value | partial slope | p-value | | |
| ≥ 2 SD | Prox-dist | 180 (36%) | 13.2 | $3 \times 10^{-4}$* | 2.94 | 0.12 | 0.0012* | 0.074 |
| | Dist-prox | 105 (21%) | -10.8 | $6 \times 10^{-8}$* | -2.76 | 0.12 | 0.0024* | 0.089 |
| | Sept-temp | 150 (30%) | 3.42 | 0.0079* | 14.1 | $9 \times 10^{-5}$* | $1 \times 10^{-4}$* | 0.069 |
| | Temp-sept | 62 (12%) | -6.25 | 0.037* | -19.6 | 0.0018* | 0.0036* | 0.049 |
| 2-4 SD | Prox-dist | 22 (48%) | 25.02 | $6 \times 10^{-4}$* | 1.83 | 0.60 | 0.001* | 0.039 |
| | Dist-prox | 7 (15%) | -19.5 | 0.0011* | -5.17 | 0.17 | 0.0047* | 0.049 |
| | Sept-temp | 9 (20%) | 2.68 | 0.32 | 37.4 | $3 \times 10^{-4}$* | $2 \times 10^{-4}$* | 0.027 |
| | Temp-sept | 8 (17%) | -8.69 | 0.11 | -36.6 | 0.0026* | 0.0035* | 0.027 |
| 4-6 SD | Prox-dist | 29 (45%) | 16.9 | $4 \times 10^{-4}$* | 2.98 | 0.22 | 0.0013* | 0.058 |
| | Dist-prox | 9 (14%) | -26.4 | $4 \times 10^{-4}$* | -13.6 | 0.014* | 0.0085* | 0.034 |
| | Sept-temp | 12 (18%) | -3.94 | 0.39 | 41.6 | 0.0015* | 0.0011* | 0.024 |
| | Temp-sept | 15 (23%) | -3.88 | 0.064 | -19.8 | $6 \times 10^{-4}$* | $7 \times 10^{-4}$* | 0.049 |
| ≥ 6 SD | Prox-dist | 129 (33%) | 10.7 | $3 \times 10^{-4}$* | 2.93 | 0.074 | 0.0015* | 0.090 |
| | Dist-prox | 89 (23%) | -8.89 | $8 \times 10^{-4}$* | -1.36 | 0.35 | 0.0021* | 0.11 |
| | Sept-temp | 129 (33%) | 3.86 | $6 \times 10^{-4}$* | 9.58 | $5 \times 10^{-5}$* | $2 \times 10^{-4}$* | 0.097 |
| | Temp-sept | 39 (10%) | -4.77 | 0.059* | -16.2 | 0.0025* | 0.0045* | 0.059 |

**Supplementary Table 13c** Ripple propagation for **rat 392.**

≥ 2 SD: number of putative events = 2828; number of propagating events = 497; percentage of propagating events = 18%; ratio for direction = 1.39; ratio for slopes = 0.490; test of proportions comparing the observed proportion of events with the 50% propagation along the proximo-distal axis expected by chance, Z = 3.3, p = $5 \times 10^{-4}$*.

2-4 SD: number of putative events = 276; number of propagating events = 46; percentage of propagating events = 17%; ratio for direction = 1.71; ratio for slopes = 0.500; test of proportions comparing the observed proportion of events with the 50% propagation along the proximo-distal axis expected by chance, Z = 1.77, p = 0.039*.

4-6 SD: number of putative events = 420; number of propagating events = 65; percentage of propagating events = 15%; ratio for direction = 1.41; ratio for slopes = 0.250; test of proportions comparing the observed proportion of events with the 50% propagation along the proximo-distal axis expected by chance, Z = 1.36, p = 0.087.

≥ 6 SD: number of putative events = 2132; number of propagating events = 386; percentage of propagating events = 18%; ratio for direction = 1.29; ratio for slopes = 0.545; test of proportions comparing the observed proportion of events with the 50% propagation along the proximo-distal axis expected by chance, Z = 2.54, p = 0.0055*.

| Amplitude group | Direction of propagation | Number of ripples | Proximo-distal | | Septo-temporal | | p-value of comparison of slopes | Resultant speed |
|---|---|---|---|---|---|---|---|---|
| | | | partial slope | p-value | partial slope | p-value | | |
| ≥ 2 SD | Prox-dist | 189 (33%) | 12.7 | $4 \times 10^{-6}$* | -0.023 | 0.98 | $2 \times 10^{-4}$* | 0.078 |
| | Dist-prox | 92 (17%) | -13.3 | $6 \times 10^{-6}$* | -1.28 | 0.39 | $5 \times 10^{-4}$* | 0.075 |
| | Sept-temp | 142 (26%) | 2.49 | 0.052 | 22.3 | $1 \times 10^{-4}$* | $3 \times 10^{-4}$* | 0.045 |
| | Temp-sept | 133 (24%) | -3.55 | 0.025* | -29.6 | $5 \times 10^{-5}$* | $1 \times 10^{-4}$* | 0.034 |
| 2-4 SD | Prox-dist | 12 (31%) | 40.1 | $3 \times 10^{-4}$* | -1.44 | 0.88 | 0.0095* | 0.025 |
| | Dist-prox | 7 (18%) | -43.04 | 0.0011* | -11.0 | 0.44 | 0.54 | 0.023 |
| | Sept-temp | 9 (23%) | -0.074 | 0.97 | 44.5 | $1 \times 10^{-4}$* | $2 \times 10^{-4}$* | 0.022 |
| | Temp-sept | 11 (28%) | -3.09 | 0.33 | -36.8 | 0.0014* | 0.0029* | 0.027 |
| 4-6 SD | Prox-dist | 20 (28%) | 21.7 | $3 \times 10^{-5}$* | -0.52 | 0.87 | 0.0015* | 0.034 |
| | Dist-prox | 8 (11%) | -29.6 | $4 \times 10^{-5}$* | -0.97 | 0.84 | 0.002* | 0.044 |
| | Sept-temp | 16 (22%) | 1.54 | 0.31 | 31.7 | $9 \times 10^{-5}$* | $2 \times 10^{-4}$* | 0.032 |
| | Temp-sept | 27 (38%) | -9.38 | 0.0025* | -39.6 | $8 \times 10^{-5}$* | $4 \times 10^{-4}$* | 0.025 |
| ≥ 6 SD | Prox-dist | 157 (35%) | 10.2 | $7 \times 10^{-6}$* | 0.076 | 0.94 | $3 \times 10^{-4}$* | 0.098 |
| | Dist-prox | 77 (17%) | -11.1 | $5 \times 10^{-6}$* | 0.64 | 0.59 | $3 \times 10^{-4}$* | 0.090 |
| | Sept-temp | 117 (26%) | 3.14 | 0.012* | 18.1 | $1 \times 10^{-4}$* | $4 \times 10^{-4}$* | 0.055 |
| | Temp-sept | 95 (21%) | -1.72 | 0.16 | -26.7 | $6 \times 10^{-5}$* | $1 \times 10^{-4}$* | 0.037 |

**Supplementary Table 13d** Ripple propagation for **rat 416.**

≥ 2 SD: number of putative events = 2520; number of propagating events = 556; percentage of propagating events = 22%; ratio for direction = 0.97; ratio for slopes = 0.636; test of proportions comparing the observed proportion of events with the 50% propagation along the proximo-distal axis expected by chance, Z = 0.25, p = 0.4.

2-4 SD: number of putative events = 202; number of propagating events = 39; percentage of propagating events = 19%; ratio for direction = 0.95; ratio for slopes = 1.500; test of proportions comparing the observed proportion of events with the 50% propagation along the proximo-distal axis expected by chance, Z = -0.16, p = 0.44.

4-6 SD: number of putative events = 407; number of propagating events = 71; percentage of propagating events = 17%; ratio for direction = 0.65; ratio for slopes = 4; test of proportions comparing the observed proportion of events with the 50% propagation along the proximo-distal axis expected by chance, Z = -1.78, p = 0.038*.

≥ 6 SD: number of putative events = 1911; number of propagating events = 446; percentage of propagating events = 23%; ratio for direction = 1.10; ratio for slopes = 0.575; test of proportions comparing the observed proportion of events with the 50% propagation along the proximo-distal axis expected by chance, Z = 1.05, p = 0.15.

| Amplitude group | Direction of propagation | Number of ripples | Proximo-distal | | Septo-temporal | | p-value of comparison of slopes | Resultant speed |
|---|---|---|---|---|---|---|---|---|
| | | | partial slope | p-value | partial slope | p-value | | |
| ≥ 2 SD | Prox-dist | 321 (31%) | 13.0 | $2 \times 10^{-4}$* | -1.64 | 0.29 | 0.0015* | 0.076 |
| | Dist-prox | 232 (23%) | -20.7 | $1 \times 10^{-4}$* | 0.36 | 0.86 | $7 \times 10^{-4}$* | 0.048 |
| | Sept-temp | 208 (20%) | 7.86 | 0.076 | 4.99 | 0.47 | 0.65 | 0.12 |
| | Temp-sept | 261 (26%) | 2.06 | 0.14 | -14.8 | $6 \times 10^{-4}$* | 0.0015* | 0.067 |
| 2-4 SD | Prox-dist | 35 (23%) | 30.05 | 0.0012* | -3.25 | 0.56 | 0.0078* | 0.033 |
| | Dist-prox | 39 (26%) | -29.6 | $2 \times 10^{-4}$* | 0.17 | 0.96 | 0.0011* | 0.034 |
| | Sept-temp | 41 (27%) | 6.26 | 0.026* | 21.7 | $9 \times 10^{-4}$* | 0.0041* | 0.044 |
| | Temp-sept | 36 (24%) | 9.38 | 0.011* | -31.1 | $4 \times 10^{-4}$* | 0.0022* | 0.031 |
| 4-6 SD | Prox-dist | 68 (30%) | 18.9 | $7 \times 10^{-4}$* | -4.38 | 0.19 | 0.0079* | 0.051 |
| | Dist-prox | 57 (25%) | -28.2 | $2 \times 10^{-4}$* | 0.54 | 0.86 | $9 \times 10^{-4}$* | 0.036 |
| | Sept-temp | 40 (18%) | 5.27 | 0.051 | 16.9 | 0.0039* | 0.015* | 0.056 |
| | Temp-sept | 62 (27%) | 4.81 | 0.0029* | -21.5 | $3 \times 10^{-5}$* | $1 \times 10^{-4}$* | 0.045 |
| ≥ 6 SD | Prox-dist | 218 (34%) | 10.8 | $4 \times 10^{-5}$* | -0.71 | 0.41 | $3 \times 10^{-4}$* | 0.093 |
| | Dist-prox | 136 (21%) | -15.6 | $1 \times 10^{-4}$* | 0.14 | 0.93 | $6 \times 10^{-4}$* | 0.064 |
| | Sept-temp | 127 (20%) | 6.23 | 0.13 | 0.59 | 0.93 | 0.78 | 0.16 |
| | Temp-sept | 163 (25%) | 0.82 | 0.37 | -14.3 | $2 \times 10^{-4}$* | $4 \times 10^{-4}$* | 0.069 |

**Supplementary Table 13e** Ripple propagation for **rat 417.**

≥ 2 SD: number of putative events = 5210; number of propagating events = 1022; percentage of propagating events = 20%; ratio for direction = 1.25; ratio for slopes = 0.637; test of proportions comparing the observed proportion of events with the 50% propagation along the proximo-distal axis expected by chance, Z = 2.63, p = 0.0043*.

2-4 SD: number of putative events = 705; number of propagating events = 151; percentage of propagating events = 21%; ratio for direction = 0.96; ratio for slopes = 1.667; test of proportions comparing the observed proportion of events with the 50% propagation along the proximo-distal axis expected by chance, Z = -0.24, p = 0.4.

4-6 SD: number of putative events = 1172; number of propagating events = 227; percentage of propagating events = 19%; ratio for direction = 1.22; ratio for slopes = 0.812; test of proportions comparing the observed proportion of events with the 50% propagation along the proximo-distal axis expected by chance, Z = 1.53, p = 0.064.

≥ 6 SD: number of putative events = 3333; number of propagating events = 644; percentage of propagating events = 19%; ratio for direction = 1.22; ratio for slopes = 0.603; test of proportions comparing the observed proportion of events with the 50% propagation along the proximo-distal axis expected by chance, Z = 2.5, p = 0.0058*.

| Amplitude group | Direction of propagation | Number of ripples | Proximo-distal | | Septo-temporal | | p-value of comparison of slopes | Resultant speed |
|---|---|---|---|---|---|---|---|---|
| | | | partial slope | p-value | partial slope | p-value | | |
| ≥ 2 SD | Prox-dist | 282 (30%) | 12.7 | $7 \times 10^{-6}$* | 1.47 | 0.31 | $2 \times 10^{-4}$* | 0.078 |
| | Dist-prox | 273 (29%) | -11.8 | $2 \times 10^{-7}$* | 1.43 | 0.068 | $4 \times 10^{-6}$* | 0.084 |
| | Sept-temp | 199 (21%) | -4.42 | 0.019* | 20.2 | $7 \times 10^{-5}$* | $3 \times 10^{-4}$* | 0.048 |
| | Temp-sept | 184 (20%) | 1.91 | 0.21 | -17.02 | $2 \times 10^{-4}$* | $3 \times 10^{-4}$* | 0.058 |
| 2-4 SD | Prox-dist | 19 (25%) | 43.2 | $2 \times 10^{-4}$* | 14.4 | 0.12 | 0.0095* | 0.022 |
| | Dist-prox | 17 (23%) | -27.3 | $7 \times 10^{-7}$* | 10.6 | 0.0014* | $1 \times 10^{-4}$* | 0.034 |
| | Sept-temp | 28 (37%) | -11.6 | 0.0093* | 34.2 | $3 \times 10^{-4}$* | 0.0027* | 0.028 |
| | Temp-sept | 11 (15%) | 9.35 | 0.0019* | -28.8 | $4 \times 10^{-5}$* | $4 \times 10^{-4}$* | 0.033 |
| 4-6 SD | Prox-dist | 59 (39%) | 15.1 | $2 \times 10^{-5}$* | 3.50 | 0.11 | $8 \times 10^{-4}$* | 0.064 |
| | Dist-prox | 22 (15%) | -28.0 | $4 \times 10^{-4}$* | 7.38 | 0.26 | 0.012* | 0.034 |
| | Sept-temp | 39 (26%) | -6.60 | 0.0085* | 27.8 | $3 \times 10^{-5}$* | $2 \times 10^{-4}$* | 0.035 |
| | Temp-sept | 30 (20%) | 3.94 | 0.0018* | -14.5 | $1 \times 10^{-5}$* | $9 \times 10^{-5}$* | 0.066 |
| ≥ 6 SD | Prox-dist | 204 (29%) | 10.3 | $8 \times 10^{-6}$* | 0.75 | 0.52 | $1 \times 10^{-4}$* | 0.097 |
| | Dist-prox | 234 (33%) | -10.4 | $8 \times 10^{-8}$* | 0.88 | 0.13 | $2 \times 10^{-6}$* | 0.096 |
| | Sept-temp | 132 (18%) | -3.16 | 0.022* | 16.9 | $3 \times 10^{-5}$* | $1 \times 10^{-4}$* | 0.058 |
| | Temp-sept | 143 (20%) | 1.17 | 0.46* | -15.8 | $4 \times 10^{-4}$* | $6 \times 10^{-4}$* | 0.063 |

**Supplementary Table 13f** Ripple propagation for **rat 432.**

≥ 2 SD: number of putative events = 3928; number of propagating events = 938; percentage of propagating events = 16%; ratio for direction = 1.48; ratio for slopes = 0.422; test of proportions comparing the observed proportion of events with the 50% propagation along the proximo-distal axis expected by chance, $Z = 5.6$, $p = 1.6 \times 10^{-8}$*.

2-4 SD: number of putative events = 428; number of propagating events = 75; percentage of propagating events = 17%; ratio for direction = 0.92; ratio for slopes = 1; test of proportions comparing the observed proportion of events with the 50% propagation along the proximo-distal axis expected by chance, $Z = -0.35$, $p = 0.36$.

4-6 SD: number of putative events = 737; number of propagating events = 150; percentage of propagating events = 20%; ratio for direction = 1.17; ratio for slopes = 0.231; test of proportions comparing the observed proportion of events with the 50% propagation along the proximo-distal axis expected by chance, Z = 2=0.98, $p = 0.16$.

≥ 6 SD: number of putative events = 2763; number of propagating events = 713; percentage of propagating events = 26%; ratio for direction = 1.59; ratio for slopes = 0.407; test of proportions comparing the observed proportion of events with the 50% propagation along the proximo-distal axis expected by chance, $Z = 6.1$, $p = 1 \times 10^{-9}$*.

| Amplitude group | Direction of propagation | Number of ripples | Proximo-distal | | Septo-temporal | | p-value of comparison of slopes | Resultant speed |
|---|---|---|---|---|---|---|---|---|
| | | | partial slope | p-value | partial slope | p-value | | |
| ≥ 2 SD | Prox-dist | 230 (26%) | 14.1 | $4 \times 10^{-7}$* | 3.85 | 0.008* | $1 \times 10^{-5}$* | 0.068 |
| | Dist-prox | 361 (40%) | -15.3 | $2 \times 10^{-7}$* | -4.34 | 0.0033* | $6 \times 10^{-6}$* | 0.063 |
| | Sept-temp | 112 (12%) | 0.12 | 0.94 | 19.4 | $2 \times 10^{-4}$* | $5 \times 10^{-5}$* | 0.052 |
| | Temp-sept | 197 (22%) | -7.35 | $4 \times 10^{-4}$* | -20.5 | $2 \times 10^{-5}$* | $7 \times 10^{-5}$* | 0.046 |
| 2-4 SD | Prox-dist | 26 (33%) | 31.8 | $3 \times 10^{-5}$* | 6.96 | 0.17 | $4 \times 10^{-4}$* | 0.031 |
| | Dist-prox | 22 (28%) | -26.2 | $3 \times 10^{-6}$* | -5.16 | 0.094 | $5 \times 10^{-5}$* | 0.037 |
| | Sept-temp | 14 (17%) | -2.11 | 0.41 | 32.9 | $1 \times 10^{-4}$* | $6 \times 10^{-5}$* | 0.030 |
| | Temp-sept | 18 (22%) | -6.53 | 0.12 | -49.4 | $1 \times 10^{-4}$* | $9 \times 10^{-5}$* | 0.020 |
| 4-6 SD | Prox-dist | 38 (34%) | 21.6 | $1 \times 10^{-4}$* | 5.62 | 0.22 | 0.0026* | 0.045 |
| | Dist-prox | 32 (29%) | -24.7 | $1 \times 10^{-5}$* | -3.98 | 0.24 | $1 \times 10^{-4}$* | 0.040 |
| | Sept-temp | 15 (14%) | -3.24 | 0.13 | 20.3 | $5 \times 10^{-4}$* | $4 \times 10^{-4}$* | 0.049 |
| | Temp-sept | 26 (23%) | -10.2 | 0.0012* | -25.8 | $1 \times 10^{-4}$* | $5 \times 10^{-4}$* | 0.036 |
| ≥ 6 SD | Prox-dist | 166 (23%) | 11.05 | $1 \times 10^{-8}$* | 2.83 | $6 \times 10^{-4}$* | $4 \times 10^{-7}$* | 0.088 |
| | Dist-prox | 307 (43%) | -14.4 | $2 \times 10^{-7}$* | -3.78 | 0.0057* | $7 \times 10^{-6}$* | 0.067 |
| | Sept-temp | 83 (12%) | 0.54 | 0.73 | 18.5 | $3 \times 10^{-4}$* | $1 \times 10^{-4}$* | 0.054 |
| | Temp-sept | 153 (22%) | -6.50 | $2 \times 10^{-5}$* | -15.7 | $2 \times 10^{-6}$* | $1 \times 10^{-5}$* | 0.059 |

**Supplementary Table 13g** Ripple propagation for **rat 441.**

≥ 2 SD: number of putative events = 3391; number of propagating events = 900; percentage of propagating events = 16%; ratio for direction = 1.93; ratio for slopes = 0.356; test of proportions comparing the observed proportion of events with the 50% propagation along the proximo-distal axis expected by chance, Z = 9.4, p = 0*.

2-4 SD: number of putative events = 323; number of propagating events = 80; percentage of propagating events = 25%; ratio for direction = 1.50; ratio for slopes = 1; test of proportions comparing the observed proportion of events with the 50% propagation along the proximo-distal axis expected by chance, Z = 1.79, p = 0.037*.

4-6 SD: number of putative events = 475; number of propagating events = 111; percentage of propagating events = 23%; ratio for direction = 1.71; ratio for slopes = 0.714; test of proportions comparing the observed proportion of events with the 50% propagation along the proximo-distal axis expected by chance, $Z = 2.75$, $p = 0.003$*.

≥ 6 SD: number of putative events = 2593; number of propagating events = 709; percentage of propagating events = 27%; ratio for direction = 2.00; ratio for slopes = 0.229; test of proportions comparing the observed proportion of events with the 50% propagation along the proximo-distal axis expected by chance, $Z = 8.9$, $p = 6 \times 10^{-17}$*.

**Supplementary Table 13** Rat-wise ripple propagation.

Statistics for ripple propagation along the proximo-distal and septo-temporal axes for ≥ 2 SD, 2-4 SD, 4-6 SD, and ≥ 6 SD for each rat separately. Partial slopes in this case are the difference in time of occurrence of ripples as a function of distance along the proximo-distal and septo-temporal axes. The unit of the partial slopes are ms/mm while that of resultant speeds are mm/ms. Most of the rats showed trends similar to the average data.

| Amplitude group | Direction of propagation | Number of ripples | Proximo-distal | | Septo-temporal | | p-value of comparison of slopes | Resultant speed |
|---|---|---|---|---|---|---|---|---|
| | | | partial slope | p-value | partial slope | p-value | | |
| ≥ 2 SD | Prox-dist | 970 (31%) | 12.7 | $6 \times 10^{-12}$* | 2.01 | 0.02* | $1 \times 10^{-8}$* | 0.078 |
| | Dist-prox | 873 (28%) | -14.3 | $4 \times 10^{-12}$* | -0.72 | 0.47 | $6 \times 10^{-9}$* | 0.07 |
| | Sept-temp | 582 (19%) | 1.60 | 0.33 | 19.4 | $7 \times 10^{-11}$* | $1 \times 10^{-7}$* | 0.051 |
| | Temp-sept | 665 (22%) | -3.79 | 0.034* | -15.3 | $2 \times 10^{-6}$* | $7 \times 10^{-4}$* | 0.064 |
| 2-4 SD | Prox-dist | 46 (25%) | 48.1 | 0.038* | 6.27 | 0.72 | 0.11 | 0.021 |
| | Dist-prox | 44 (24%) | -34.1 | $1 \times 10^{-8}$* | 0.58 | 0.87 | $3 \times 10^{-6}$* | 0.029 |
| | Sept-temp | 56 (30%) | 4.53 | 0.26 | 45.7 | $2 \times 10^{-10}$* | $4 \times 10^{-7}$* | 0.022 |
| | Temp-sept | 38 (21%) | 1.54 | 0.75 | -36.6 | $2 \times 10^{-5}$* | $4 \times 10^{-4}$* | 0.027 |
| 4-6 SD | Prox-dist | 143 (33%) | 16.3 | 0.0019* | -1.54 | 0.46 | 0.0086* | 0.061 |
| | Dist-prox | 90 (21%) | -22.1 | $3 \times 10^{-7}$* | -4.27 | 0.12 | $3 \times 10^{-4}$* | 0.044 |
| | Sept-temp | 91 (21%) | 1.01 | 0.76 | 30.4 | $2 \times 10^{-8}$* | $5 \times 10^{-6}$* | 0.033 |
| | Temp-sept | 105 (25%) | -8.66 | 0.0036* | -19.3 | $6 \times 10^{-5}$* | 0.026* | 0.047 |
| ≥ 6 SD | Prox-dist | 781 (31%) | 10.6 | $1 \times 10^{-12}$* | 2.59 | $4 \times 10^{-4}$* | $4 \times 10^{-8}$* | 0.092 |
| | Dist-prox | 739 (30%) | -12.2 | $1 \times 10^{-11}$* | 0.53 | 0.56 | $1 \times 10^{-8}$* | 0.082 |
| | Sept-temp | 435 (18%) | 1.55 | 0.27 | 15.8 | $2 \times 10^{-10}$* | $4 \times 10^{-7}$* | 0.063 |
| | Temp-sept | 522 (21%) | -3.42 | 0.03* | -14.2 | $8 \times 10^{-7}$* | $4 \times 10^{-4}$* | 0.068 |

**Supplementary Table 14** Ripple propagation for comparable spatial spreads.

Statistics for ripple propagation along the proximo-distal and septo-temporal axes for ≥ 2 SD, 2-4 SD, 4-6 SD, and ≥ 6 SD after controlling for differences in spatial spreads of tetrodes along the proximo-distal and septo-temporal axes. Partial slopes in this case are the difference in time of occurrence of ripples as a function of distance along the proximo-distal and septo-temporal axes. The unit of the partial slopes are ms/mm while that of resultant speeds are mm/ms.

≥ 2 SD: number of putative events = 19253; number of propagating events = 3090; percentage of propagating events = 16%; ratio for direction = 1.48; ratio for slopes = 0.513; test of proportions comparing the observed proportion of events with the 50% propagation along the proximo-distal axis expected by chance, Z = 11, p = 0*.

2-4 SD: number of putative events = 1374; number of propagating events = 184; percentage of propagating events = 13%; ratio for direction = 0.96; ratio for slopes = 0.857; test of proportions comparing the observed proportion of events with the 50% propagation along the proximo-distal axis expected by chance, $Z = -0.29$, $p = 0.38$.

4-6 SD: number of putative events = 3052; number of propagating events = 429; percentage of propagating events = 14%; ratio for direction = 1.19; ratio for slopes = 0.5; test of proportions comparing the observed proportion of events with the 50% propagation along the proximo-distal axis expected by chance, $Z = 1.79$, $p = 0.037$*.

≥ 6 SD: number of putative events = 14827; number of propagating events = 2477; percentage of propagating events = 17%; ratio for direction = 1.59; ratio for slopes = 0.497; test of proportions comparing the observed proportion of events with the 50% propagation along the proximo-distal axis expected by chance, $Z = 11$, $p = 0$*.

Consistent with the original data, the bias for propagation along the proximo-distal axis was higher for subgroups with higher amplitude on the reference tetrode, and the speeds remained similar to those in the original data for all amplitude groups.

#### Supplementary Figure 1

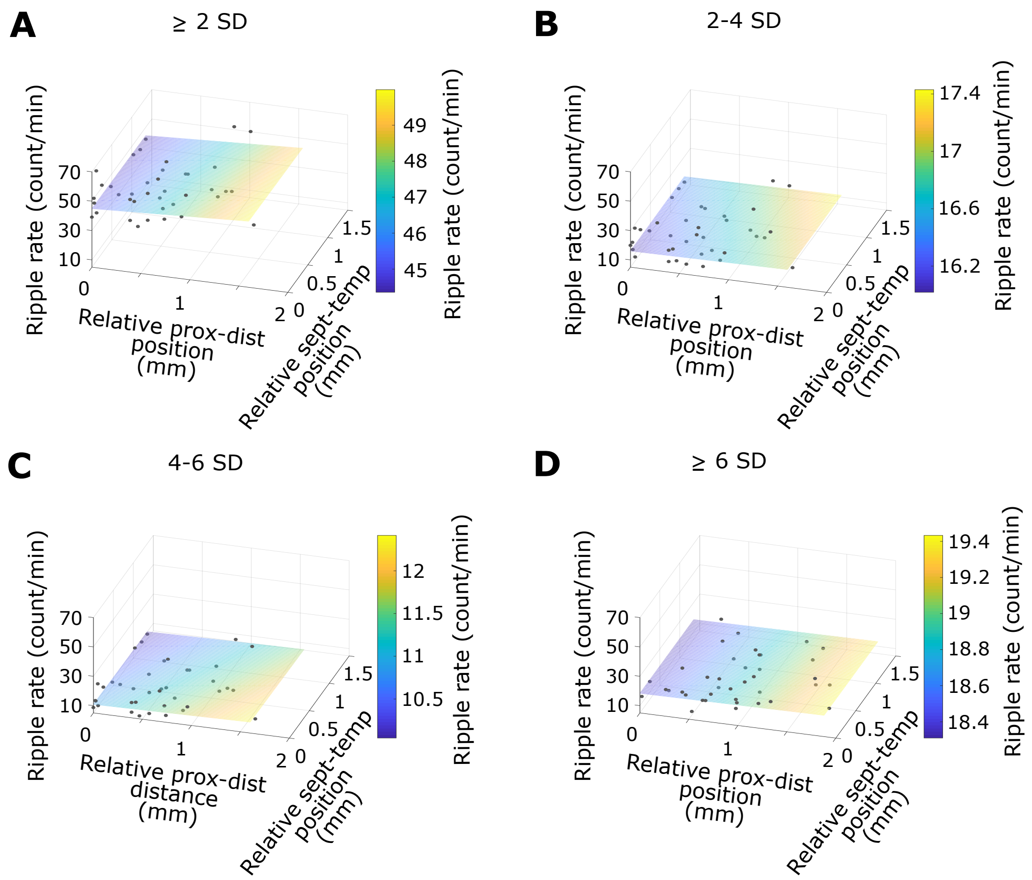


**Supplementary Figure 1** Rate of ripple occurrence using comparable spatial spreads along the proximo-distal and septo-temporal axes.

Rate of ripple occurrence as a function of relative position along the proximo-distal and septo-temporal axes for all rats with the 2D fit (plane) for ≥ 2 SD (A), 2-4 SD (B), 4-6 SD (C), and ≥ 6 SD (D).

### Supplementary Figure 2

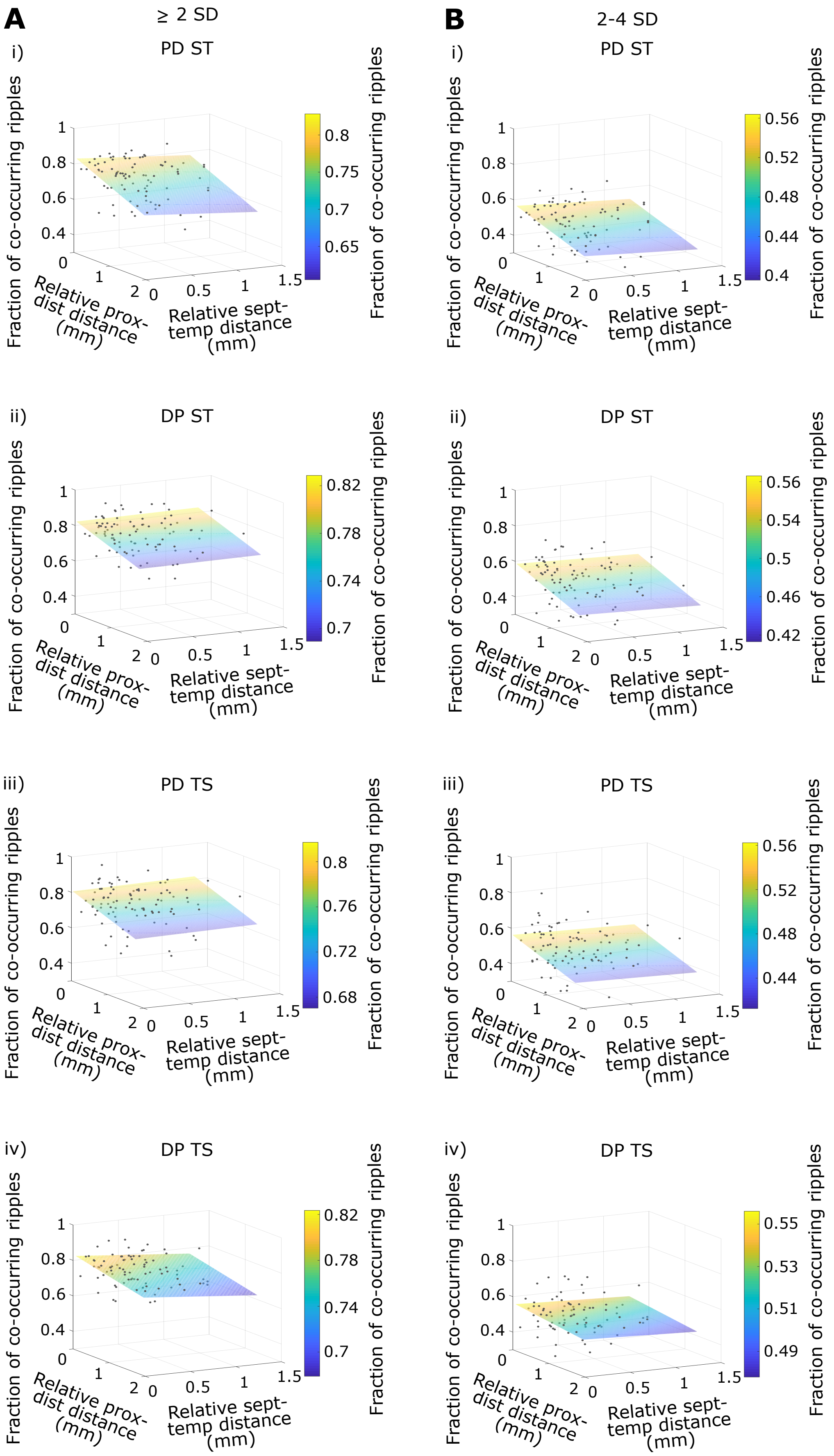

**Supplementary Figure 2 (contd.) (see next page for figure legend)**

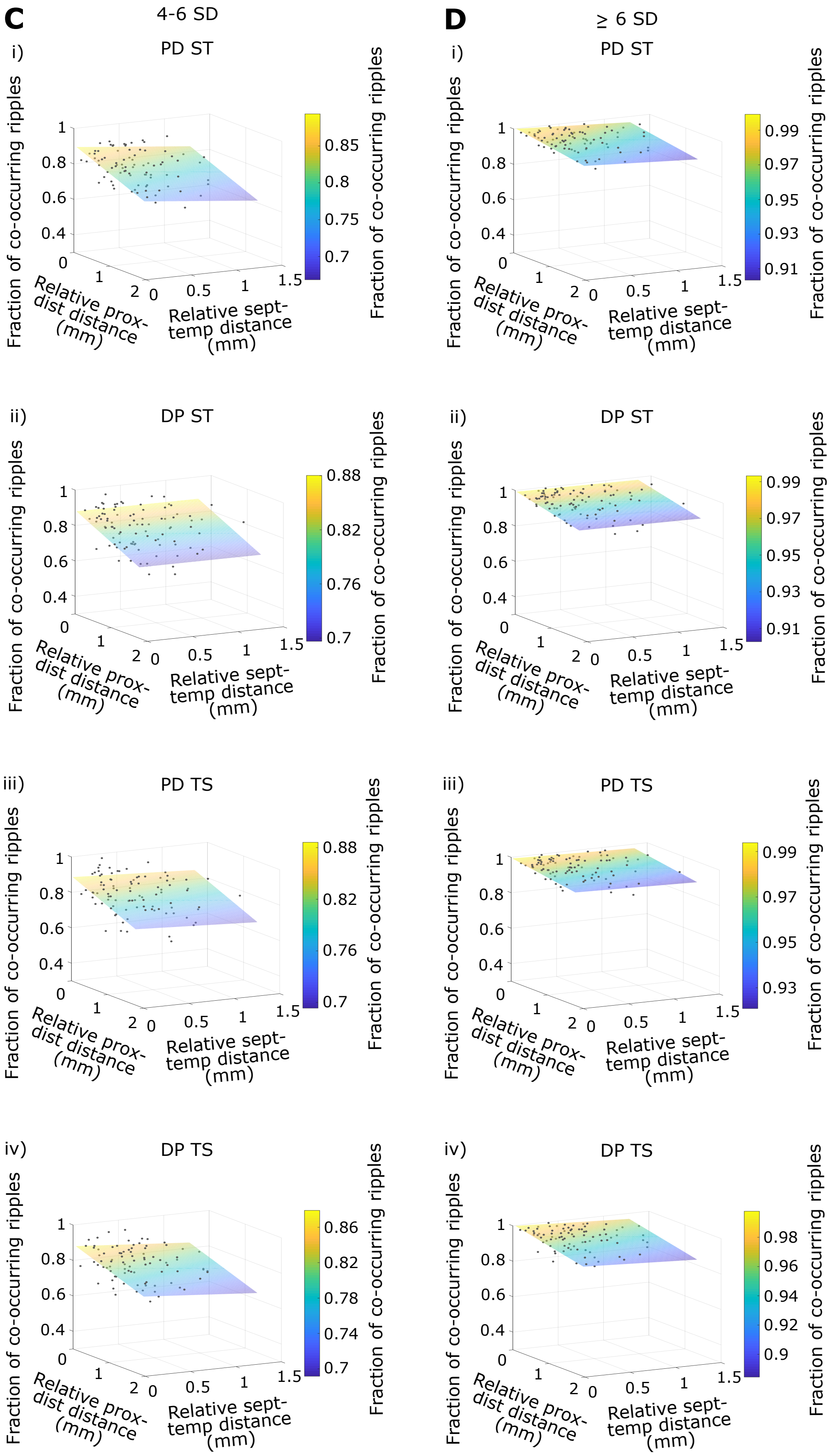

**Supplementary Figure 2** Ripple co-occurrence accounting for the direction from the reference to the referred tetrode.

(A) Fraction of co-occurring ripples for tetrode pairs having (i) reference tetrode more proximal and septal to referred tetrode (ii) reference tetrode more distal and septal to referred tetrode (iii) reference tetrode more proximal and temporal to referred tetrode (iv) reference tetrode more distal and temporal to referred tetrode with the 2D fit (plane) obtained from multiple linear regression analysis.

(B)-(D) Same as (A) but for 2-4 SD, 4-6 SD, and ≥ 6 SD.

#### Supplementary Figure 3

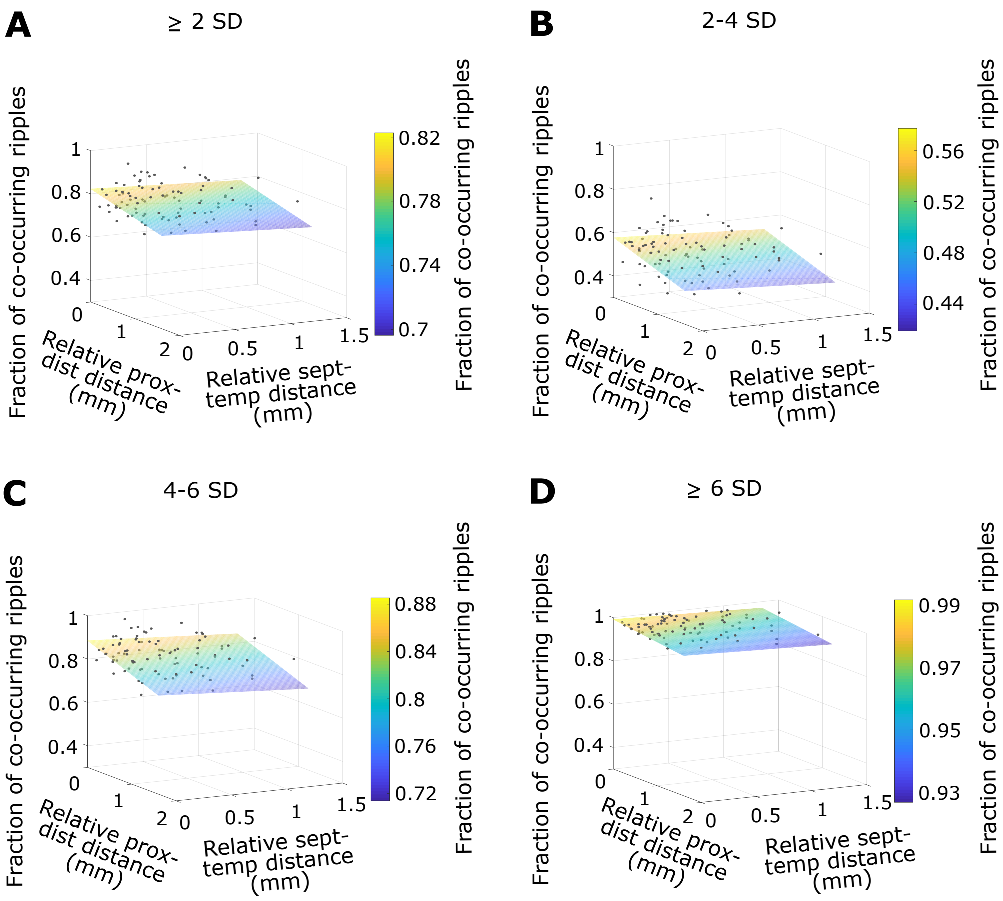


**Supplementary Figure 3** Ripple co-occurrence using comparable spatial spreads along the proximo-distal and septo-temporal axes.

Fraction of co-occurring ripples as a function of relative distance along the proximo-distal and septo-temporal axes for all rats with the 2D fit (plane) for ≥ 2 SD (A), 2-4 SD (B), 4-6 SD (C), and ≥ 6 SD (D).

### Supplementary Figure 4

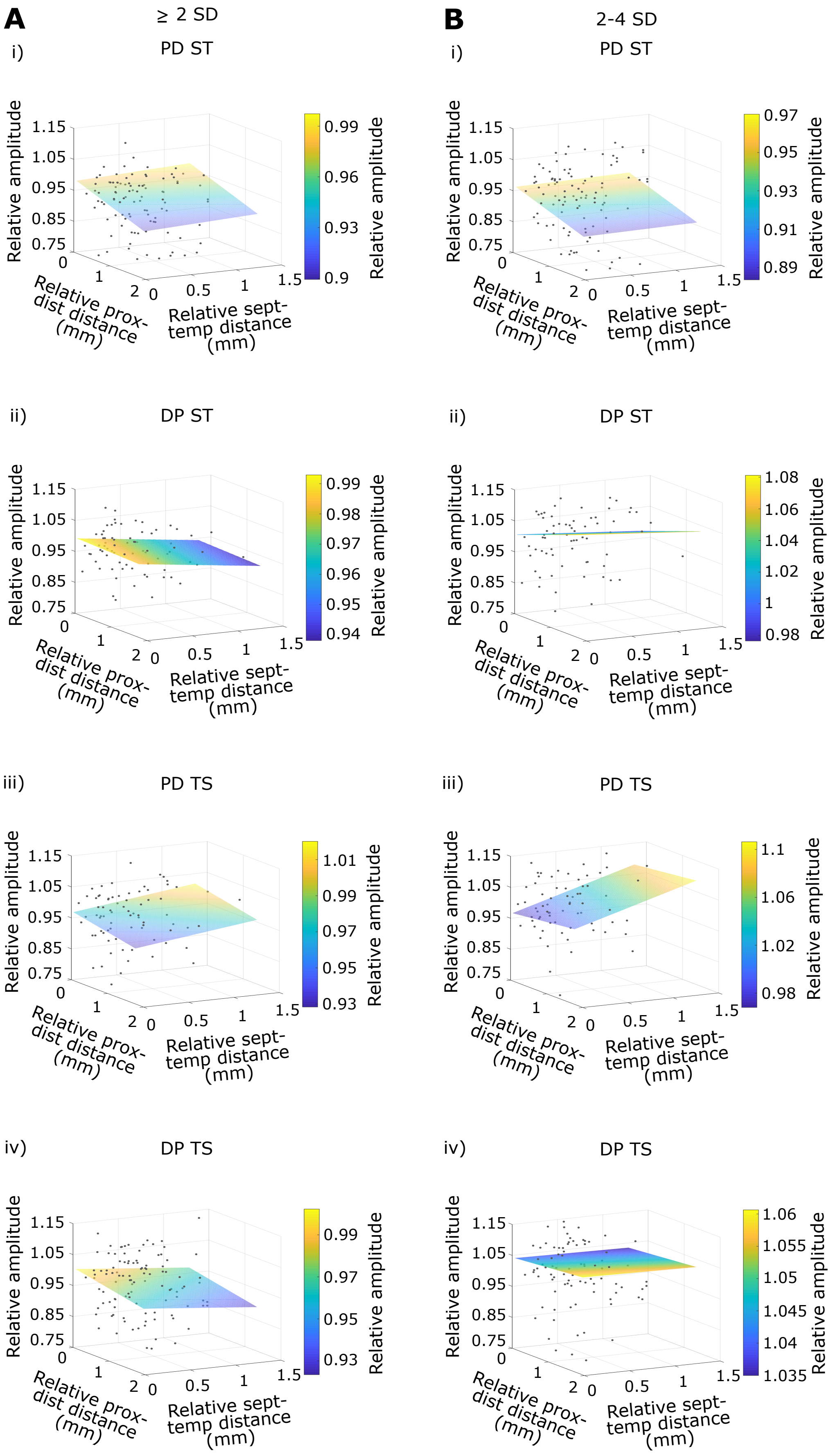

#### Supplementary Figure 4 (contd.) (see next page for figure legend)

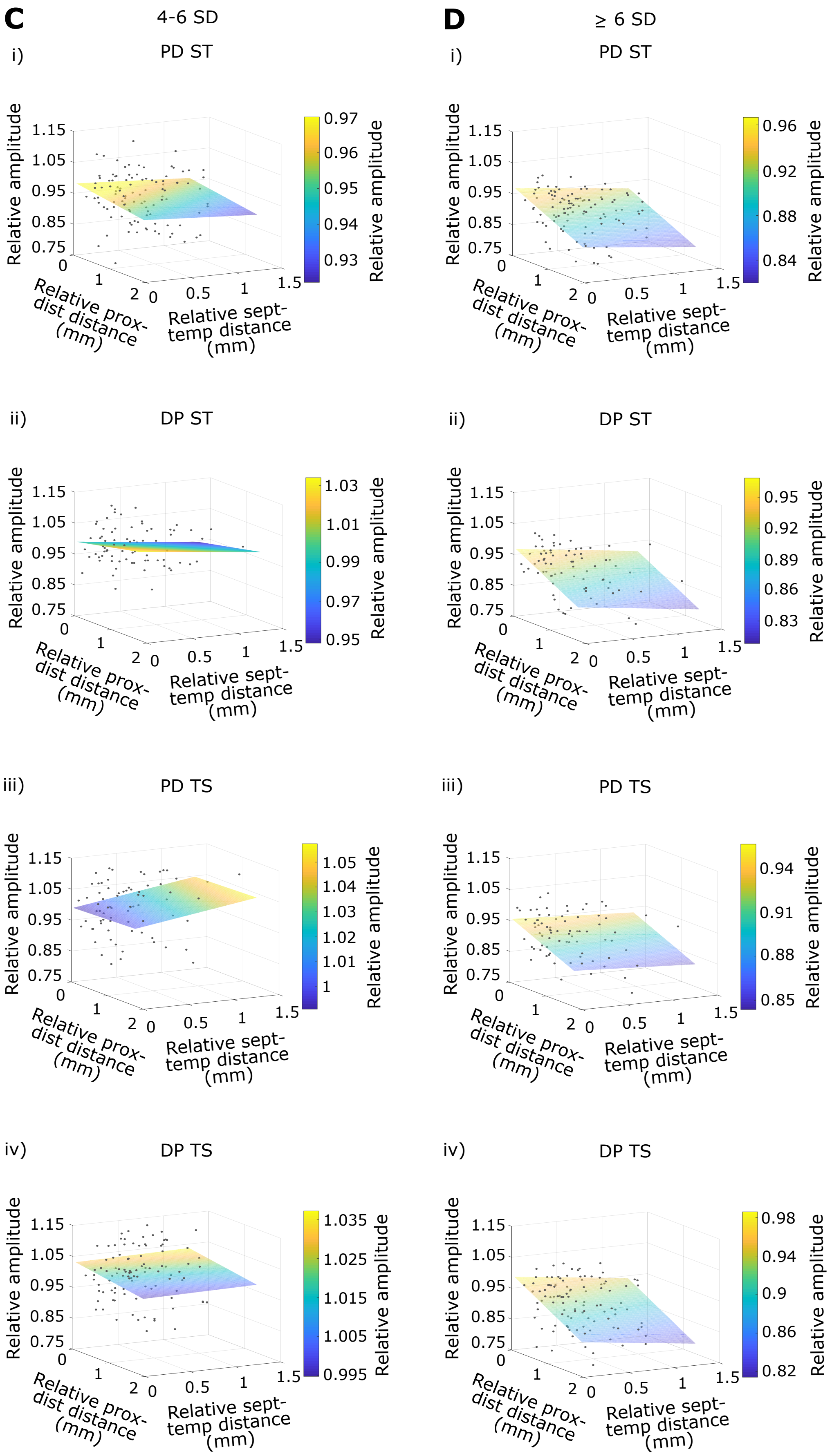

**Supplementary Figure 4** Relative ripple amplitude accounting for the direction from the reference to the referred tetrode.

Figure organization is the same as that in Supplementary Figure 2, but for relative ripple amplitude.

### Supplementary Figure 5

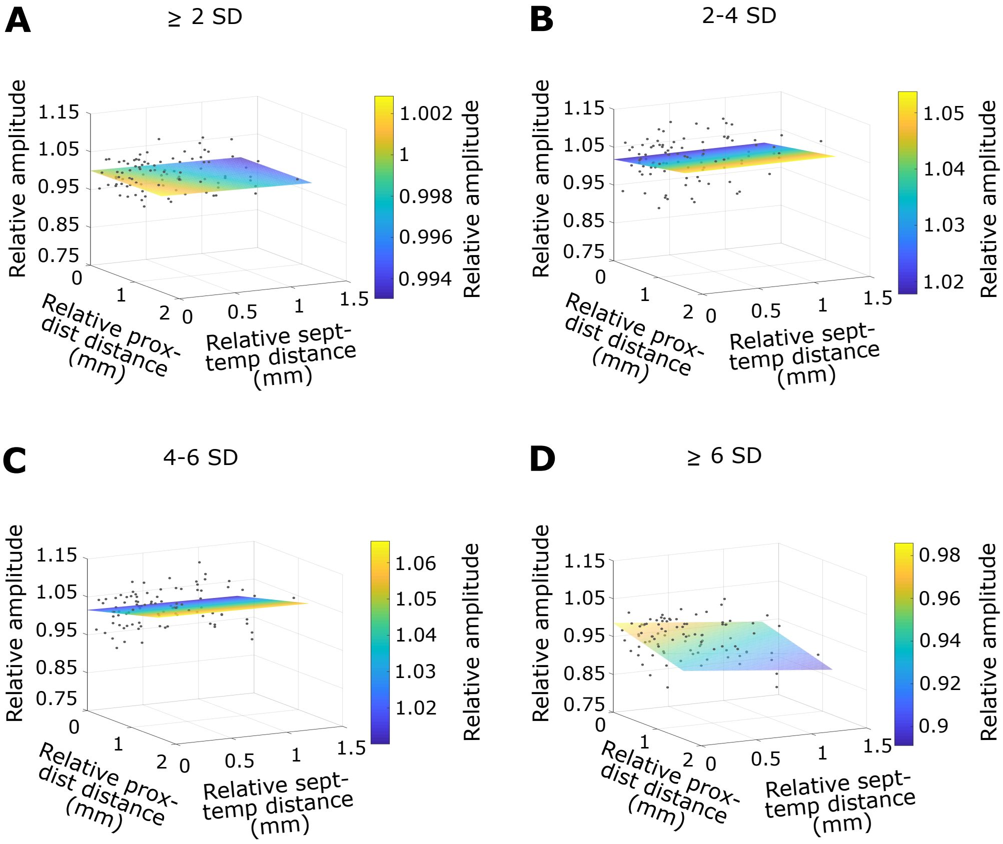


**Supplementary Figure 5** Relative ripple amplitude using comparable spatial spreads along the proximo-distal and septo-temporal axes.

Relative ripple amplitude as a function of relative distance along the proximo-distal and septo-temporal axes for all rats with the 2D fit (plane) for ≥ 2 SD (A), 2-4 SD (B), 4-6 SD (C), and ≥ 6 SD (D).

#### Supplementary Figure 6

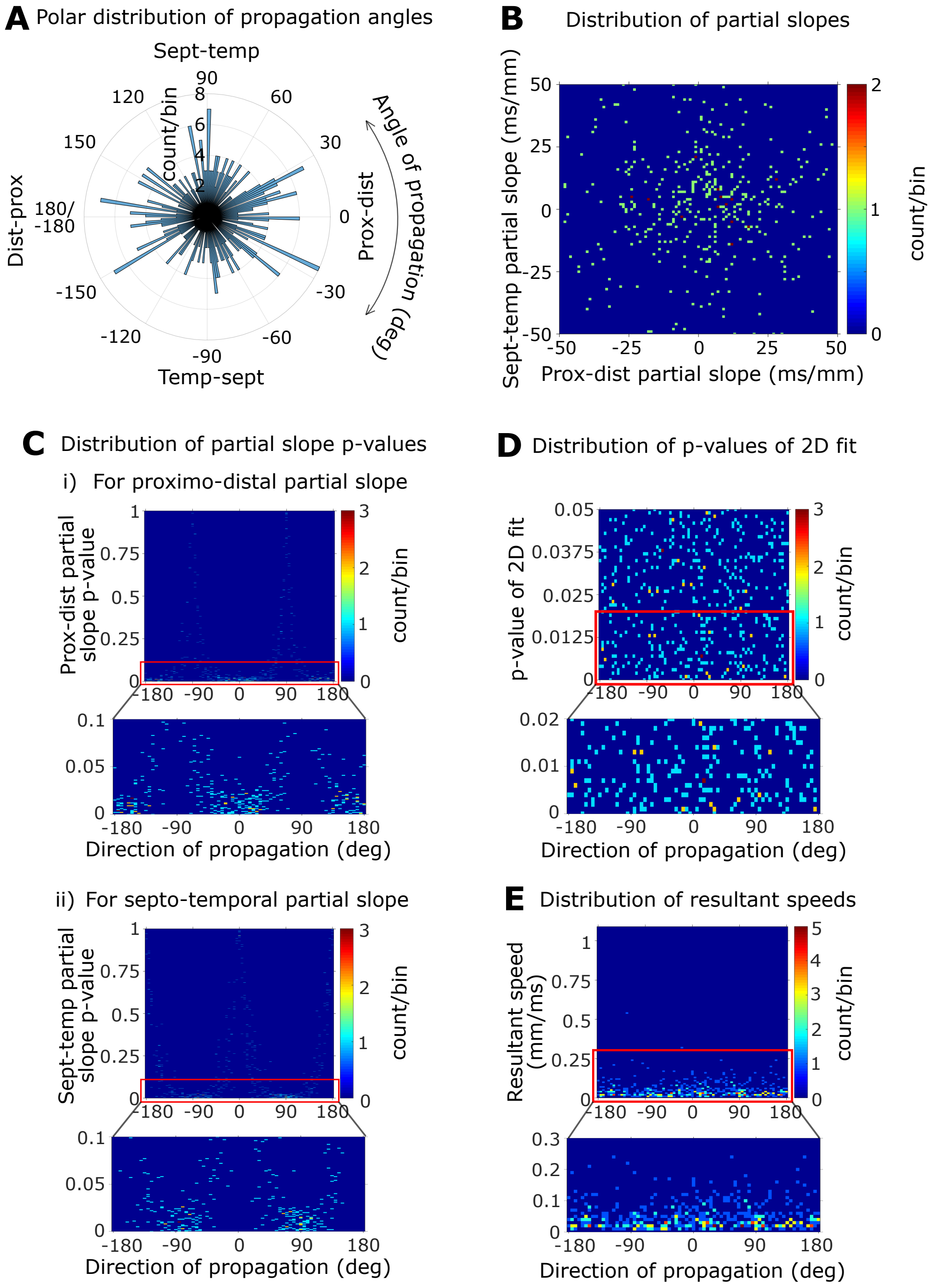


**Supplementary Figure 6** Ripple propagation for 2-4 SD.

(A) Polar distribution of the angles of propagation. The following angles correspond to the given directions: proximo-distal direction, 0°; disto-proximal direction, 180° or -180°; septo-temporal direction, 90°; temporo-septal direction, -90°.

(B) Distribution of proximo-distal vs. septo-temporal partial slopes obtained from multiple linear regression analysis of individual propagating events.

(C) Distributions of the p-values of partial slopes vs. direction (angle) of propagation for all propagating events for proximo-distal partial slopes (i) and septo-temporal partial slopes (ii). Note that the p-values are close to 0 about 0°, -180°, and 180° and close to 1 about -90° and 90° in (i), and vice versa for the p-values in (ii).

(D) Distribution of the p-values of the multiple linear regression models vs. direction of propagation for all propagating events.

(E) Distribution of the resultant speed (obtained from vector analysis) vs. direction of propagation for all propagating events.

### Supplementary Figure 7

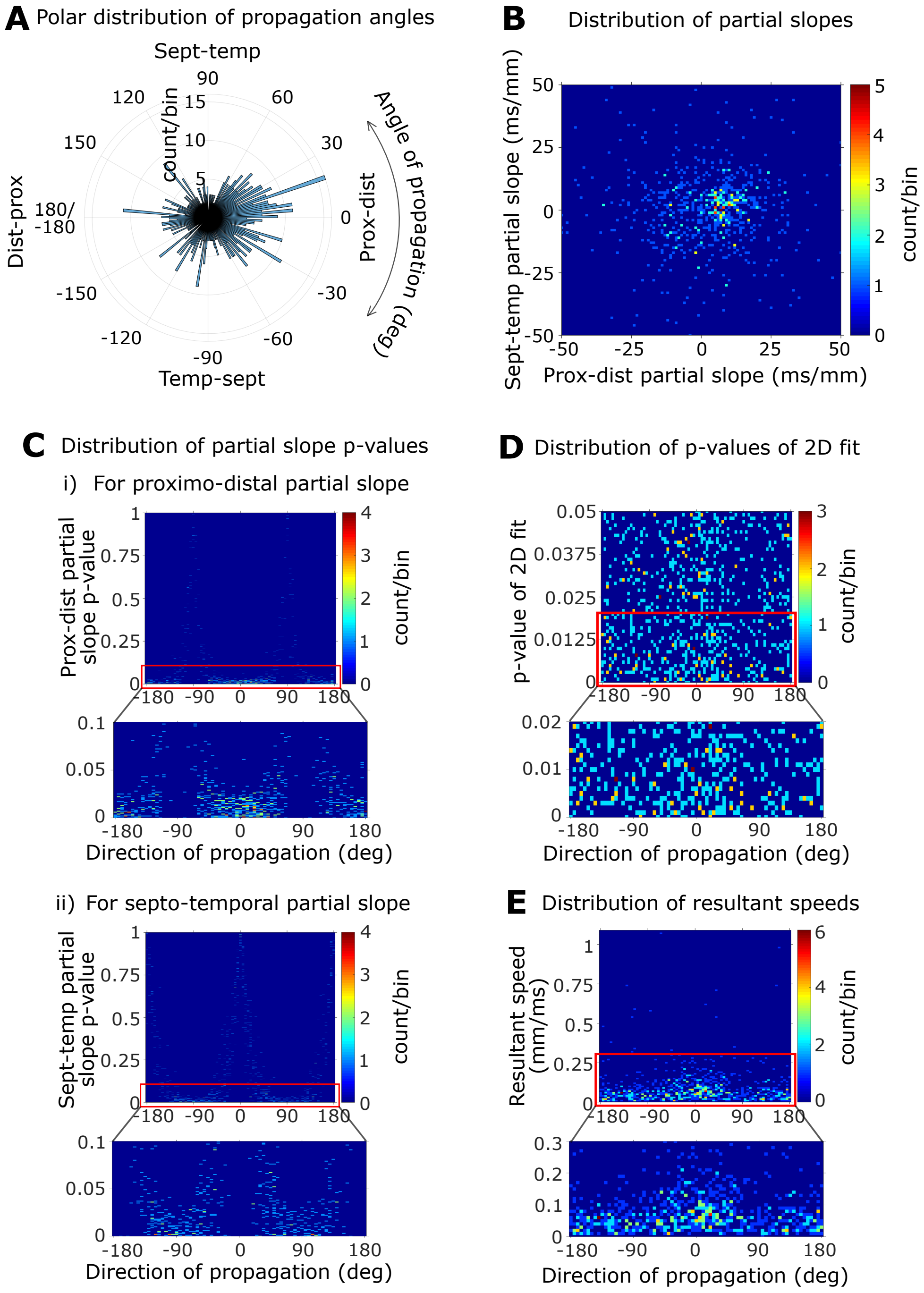


**Supplementary Figure 7** Ripple propagation for 4-6 SD.

See Supplementary Figure 6 legend for description.

### Supplementary Figure 8

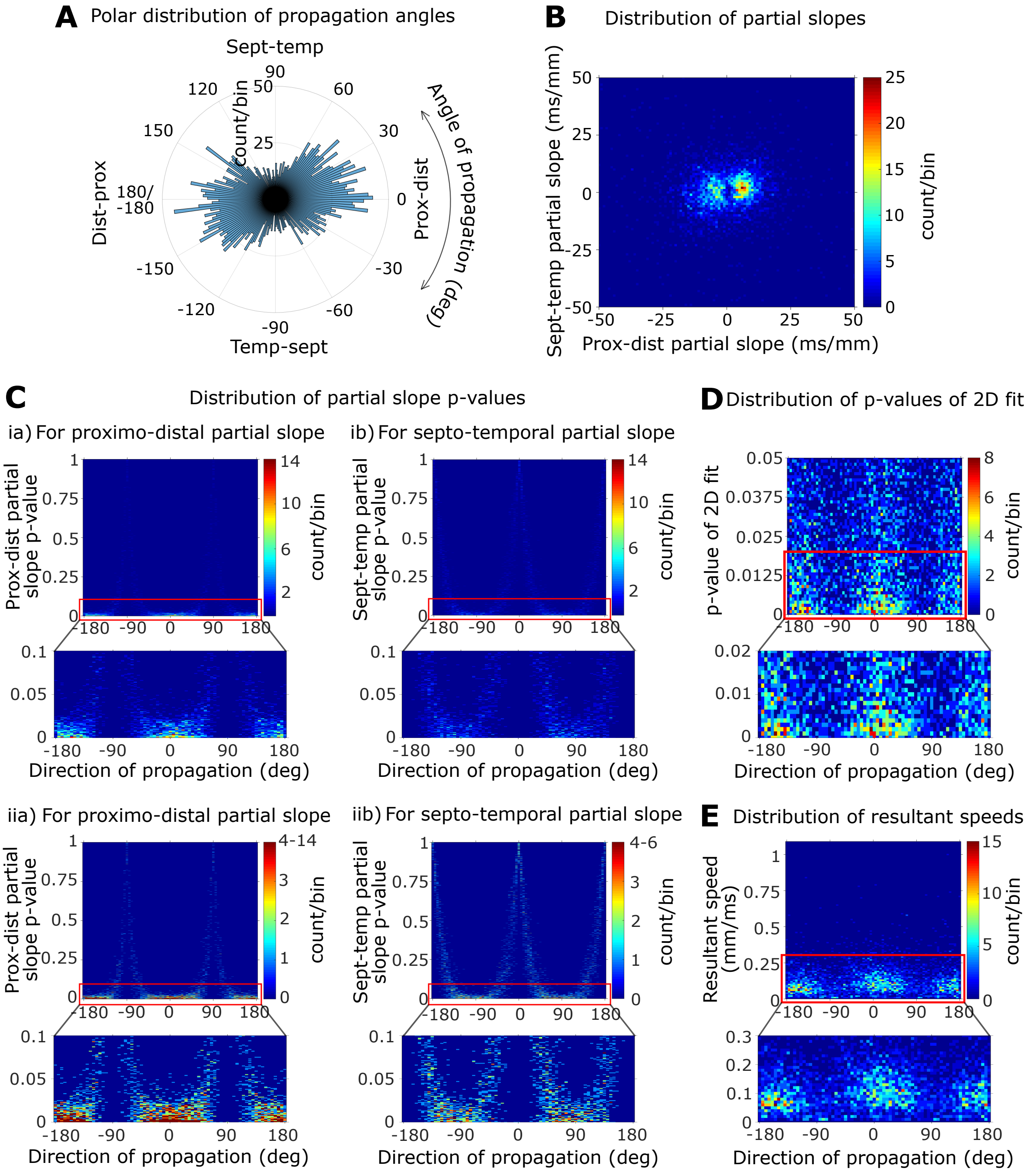


**Supplementary Figure 8** Ripple propagation for ≥ 6 SD.

See Supplementary Figure 6 legend for description. Note the clear preference for the proximo-distal axis over the septo-temporal axis in (A). Note the bimodal nature of the distribution of the proximo-distal partial slopes but not the septo-temporal partial slopes in (C). In (D), panels (i) and (ii) are the same plots, however, the color scheme of (ii) has an upper limit of 4 to facilitate visualization of the pattern – note that the p-values are close to 0 about 0°, -180°, and 180° and close to 1 about -90° and 90° in (iia) and vice versa for the p-values in (iib).

### Supplementary Figure 9 (see next page for figure legend)

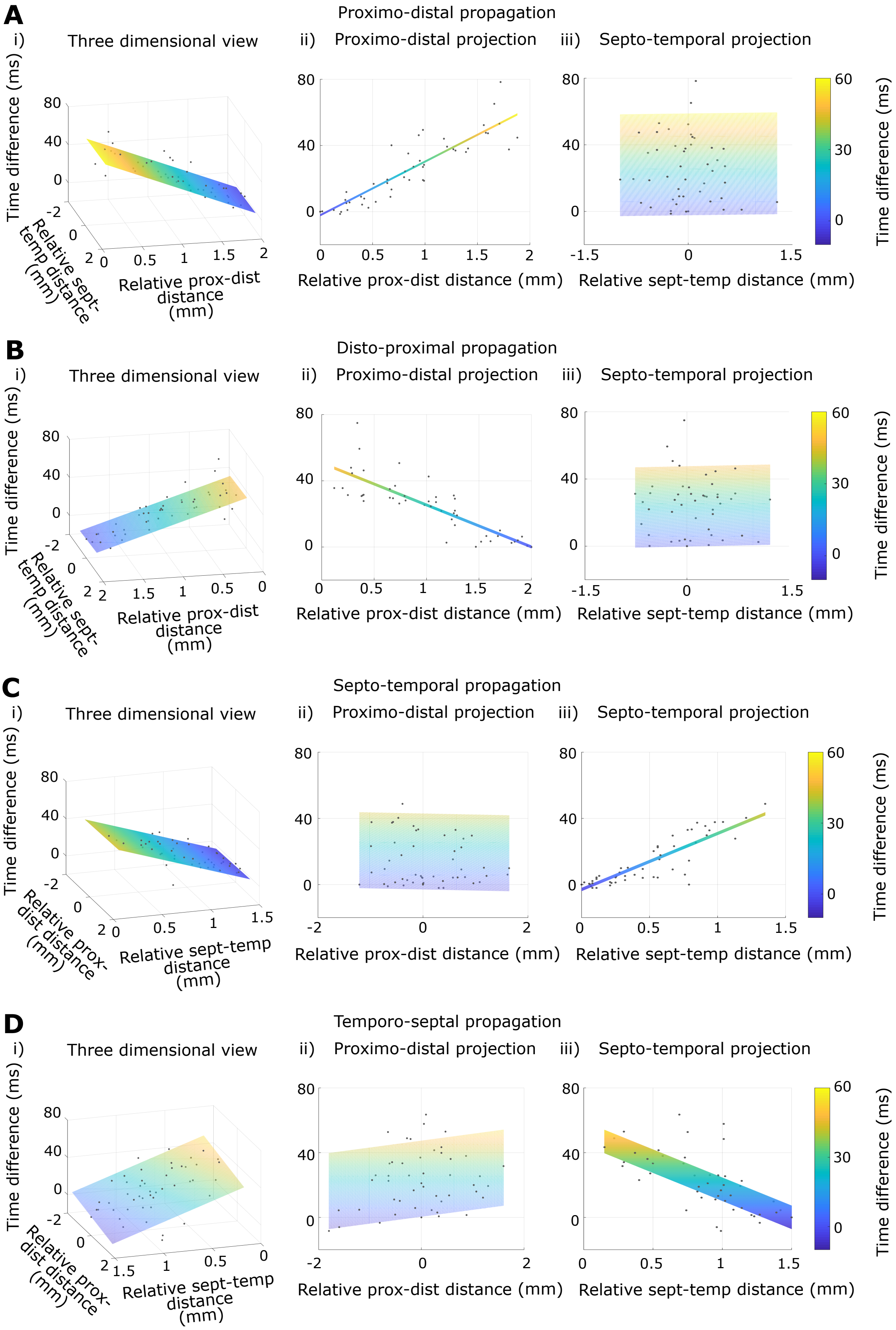

**Supplementary Figure 9** Ripple propagation speed for 2-4 SD.

(A)-(D) Distributions of relative time differences vs. relative distances along the proximo-distal and septo-temporal axes after classification of events into the four directions of propagation show clear narrow, linear trends along the expected direction of propagation. Column (i) shows a 3D (X-Y-Z) view, (ii) shows a proximo-distal projection (X-Z view), and (iii) shows a septo-temporal projection (Y-Z view) of column (i). Orientation of plots for propagation along the proximo-distal axis (Ai and Bi) are different from those along the septo-temporal axis (Ci and Di) for visualization purposes.

#### Supplementary Figure 10 (see next page for figure legend)

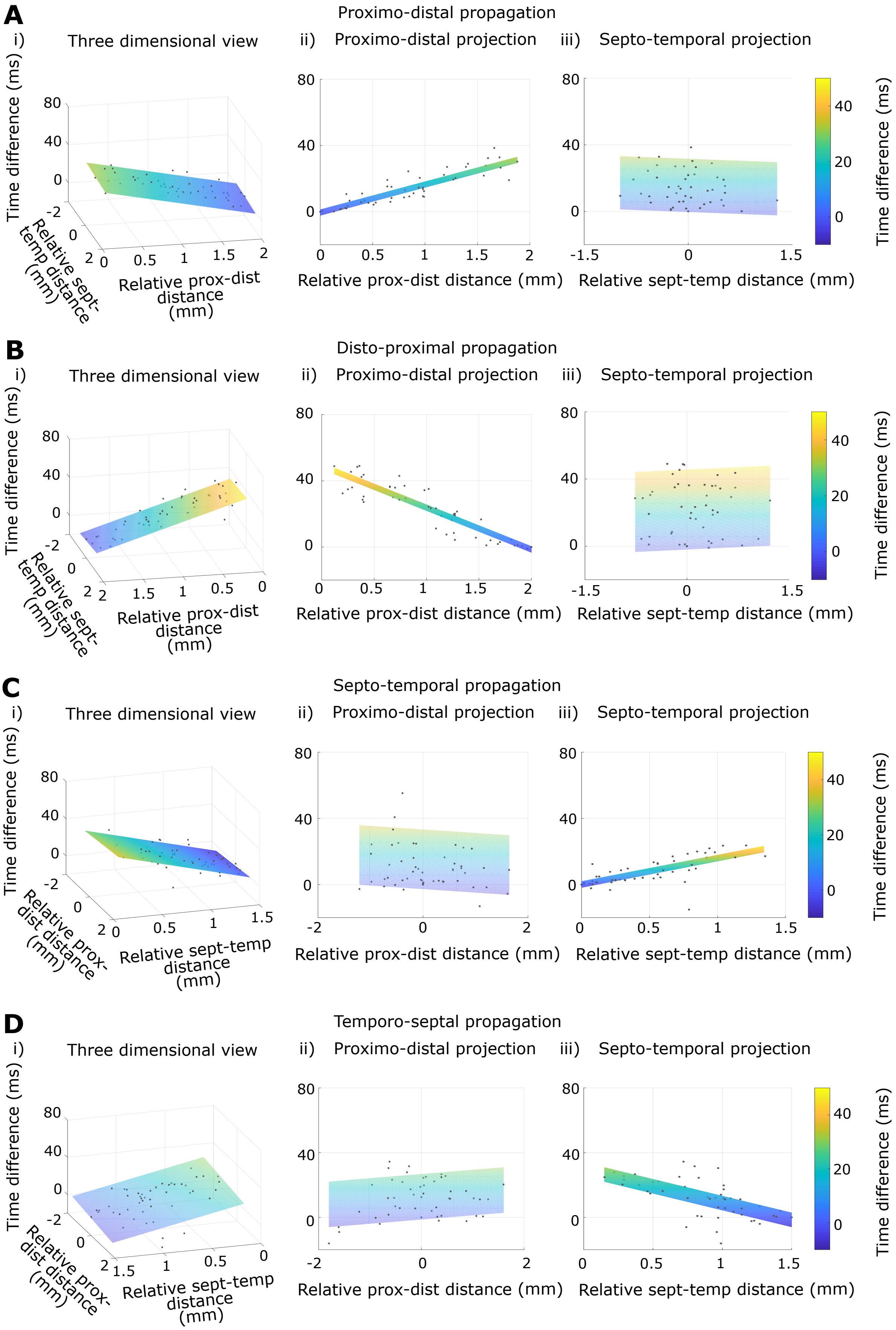

**Supplementary Figure 10** Ripple propagation speed for 4-6 SD.

See Supplementary Figure 9 legend for description.

#### Supplementary Figure 11 (see next page for figure legend)

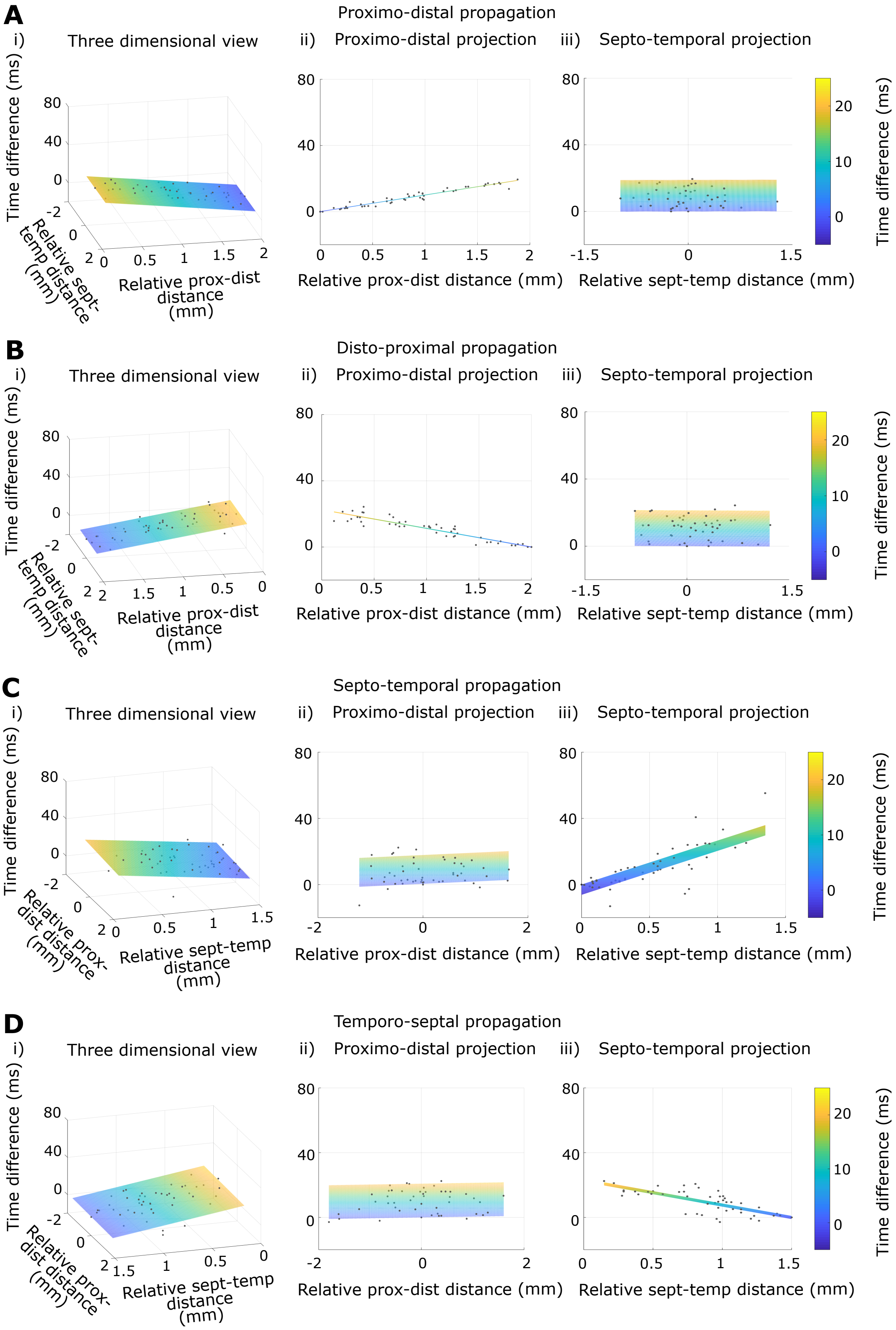

**Supplementary Figure 11** Ripple propagation speed for ≥ 6 SD.

See Supplementary Figure 9 legend for description.

#### Supplementary Figure 12 (see next page for figure legend)

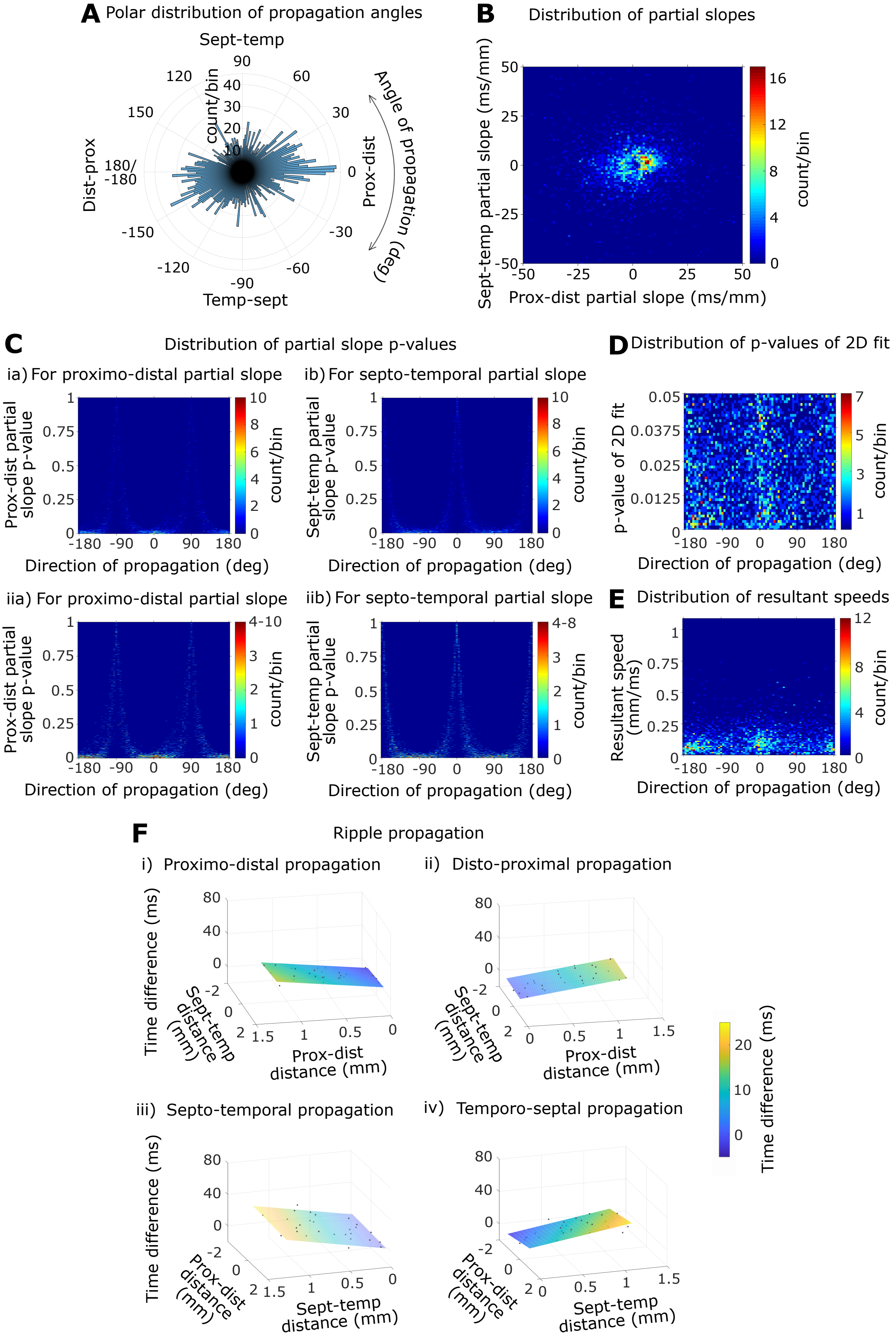

**Supplementary Figure 12** Ripple propagation using comparable spatial spreads along the proximo-distal and septo-temporal axes for ≥ 2 SD.

(A) Polar distribution of the angles of propagation. Note the clear preference for the proximo-distal axis.

(B) Distribution of proximo-distal vs. septo-temporal partial slopes obtained from multiple linear regression analysis of individual propagating events. Note the bimodal nature of the distribution of proximo-distal partial slopes but not the septo-temporal partial slopes.

(C) Distributions of the p-values of partial slopes vs. direction (angle) of propagation for all propagating events. Panels (i) and (ii) are the same plots, however, the color scheme of (ii) has an upper limit of 4 to facilitate visualization of the pattern – note that the p-values are close to 0 about 0°, -180°, and 180° and close to 1 about -90° and 90° in (iia), and vice versa for the p-values in (iib). Note the relatively high number of events about 0°, -180°, and 180° in all panels.

(D) Distribution of the p-value of the multiple linear regression models vs. direction (angle) of propagation for all propagating events.

(E) Distribution of the resultant speed (obtained from vector analysis) vs. direction (angle) of propagation for all propagating events.

(F) Distributions of relative time differences vs. relative distances along the proximo-distal and septo-temporal axes after classification of events into the four directions of propagation (i-iv) show clear narrow, linear trends along the expected direction of propagation. Orientation of plots for propagation along the proximo-distal axis (i-ii) are different from those along the septo-temporal axis (iii-iv) for visualization purposes.

#### Supplementary Figure 13 (see next page for figure legend)

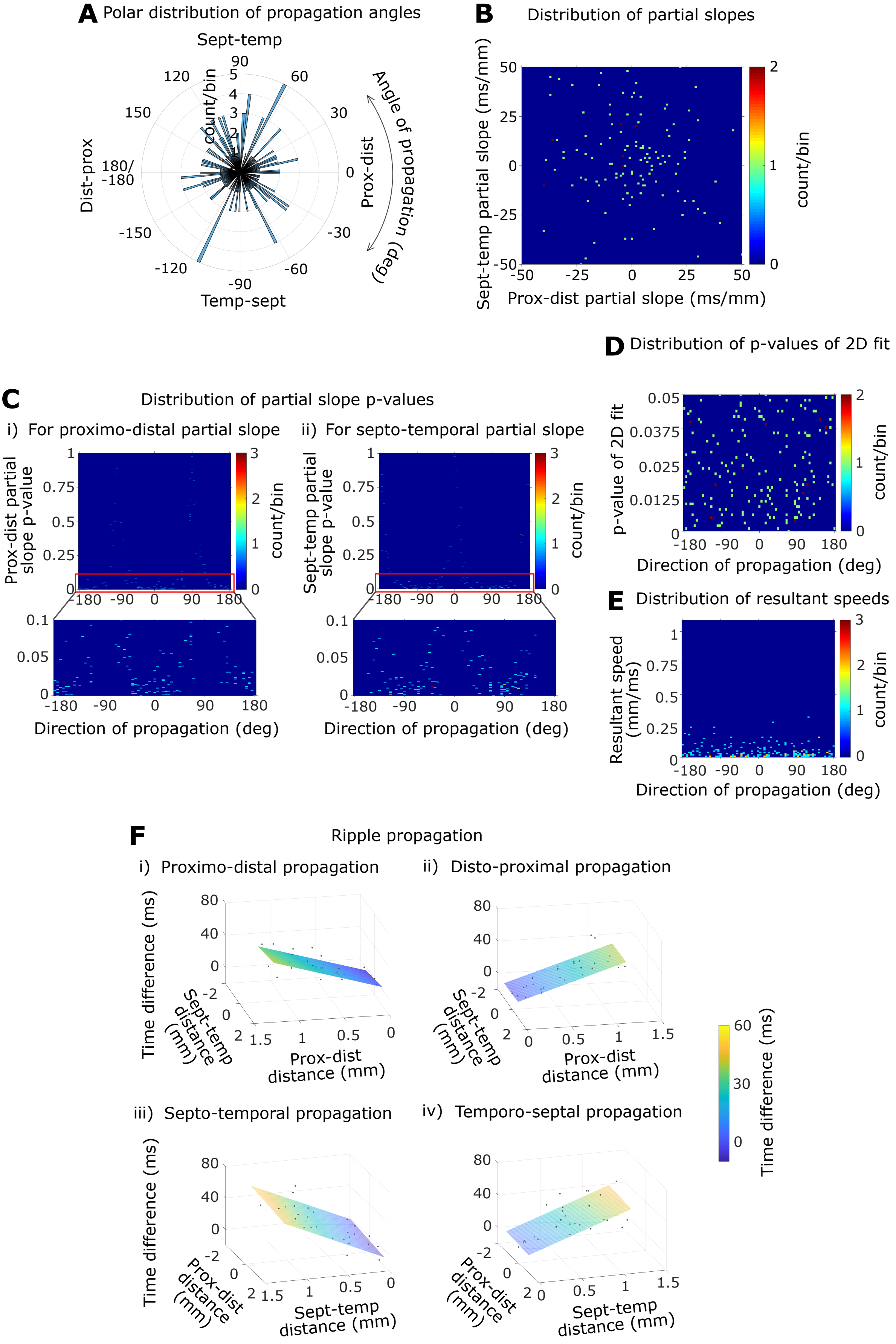

**Supplementary Figure 13** Ripple propagation using comparable spatial spreads along the proximo-distal and septo-temporal axes for 2-4 SD.

(A) Polar distribution of the angles of propagation.

(B) Distribution of proximo-distal vs. septo-temporal partial slopes obtained from multiple linear regression analysis of individual propagating events.

(C) Distributions of the p-values of partial slopes vs. direction (angle) of propagation for all propagating events. Note that the p-values are close to 0 about 0°, -180°, and 180° and close to 1 about -90° and 90° in (i), and vice versa for the p-values in (ii).

(D) Distribution of the p-value of the multiple linear regression models vs. direction (angle) of propagation for all propagating events.

(E) Distribution of the resultant speed (obtained from vector analysis) vs. direction (angle) of propagation for all propagating events.

(A) Distributions of relative time differences vs. relative distances along the proximo-distal and septo-temporal axes after classification of events into the four directions of propagation (i-iv) show clear narrow, linear trends along the expected direction of propagation. Orientation of plots for propagation along the proximo-distal axis (i-ii) are different from those along the septo-temporal axis (iii-iv) for visualization purposes.

### Supplementary Figure 14 (see next page for figure legend)

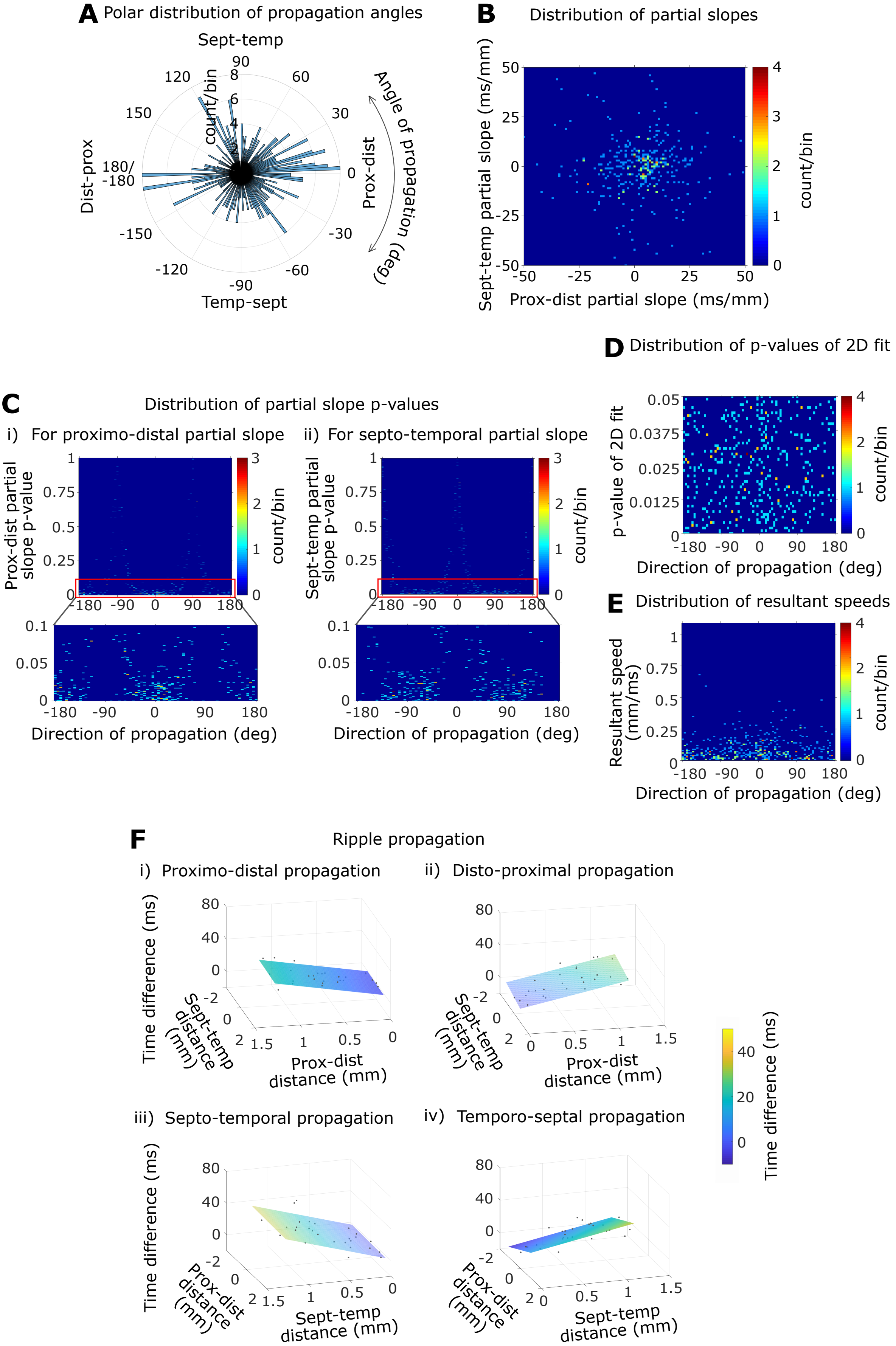

**Supplementary Figure 14** Ripple propagation using comparable spatial spreads along the proximo-distal and septo-temporal axes for 4-6 SD.

See Supplementary Figure 13 legend for description.

#### Supplementary Figure 15 (see next page for figure legend)

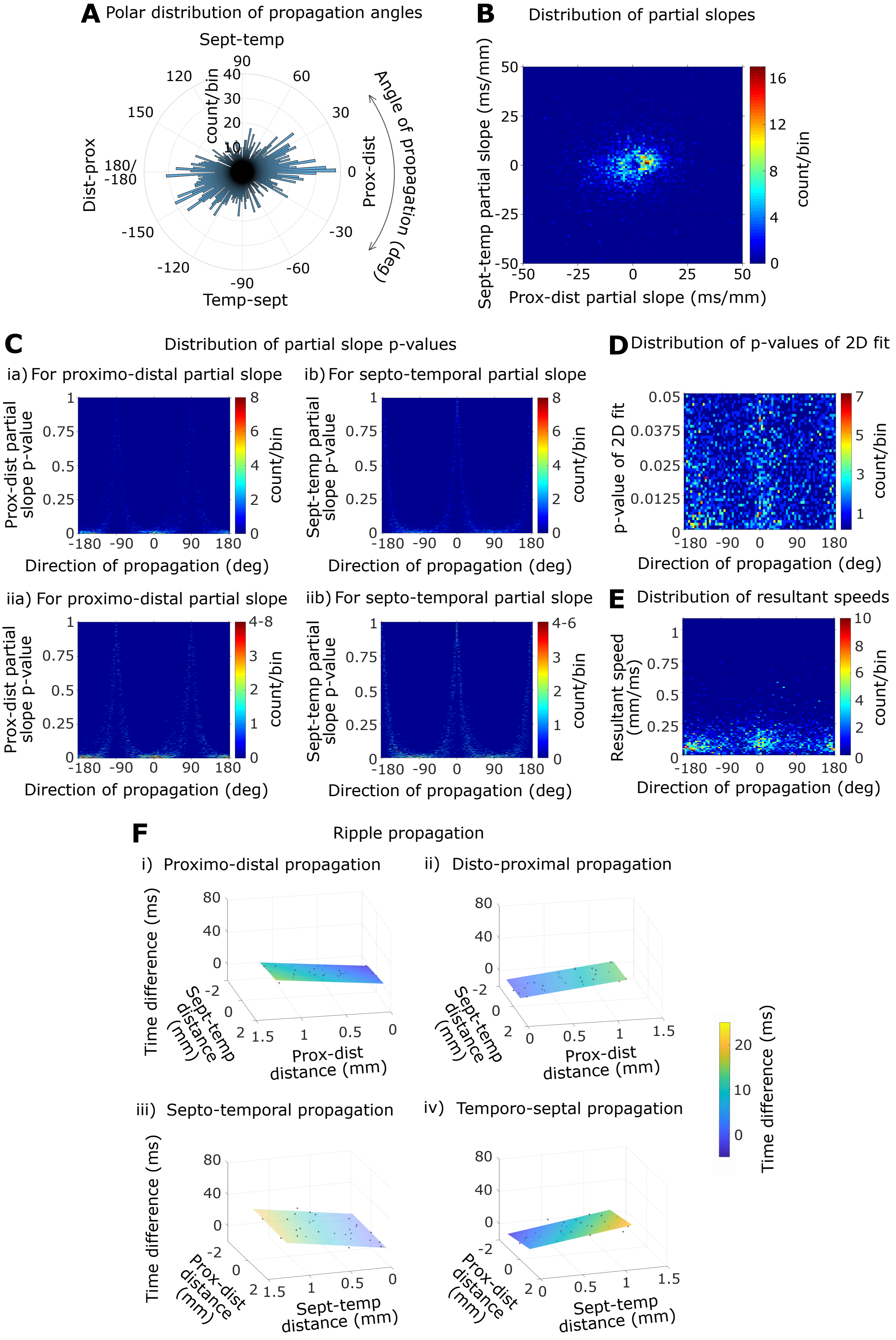

**Supplementary Figure 15** Ripple propagation using comparable spatial spreads along the proximo-distal and septo-temporal axes for ≥ 6 SD.

See Supplementary Figure 12 legend for description. Note the clear preference for the proximo-distal axis over the septo-temporal axis in (A). Note the bimodal nature of the distribution of the proximo-distal partial slopes but not the septo-temporal partial slopes in (B).

### Supplementary Figure 16

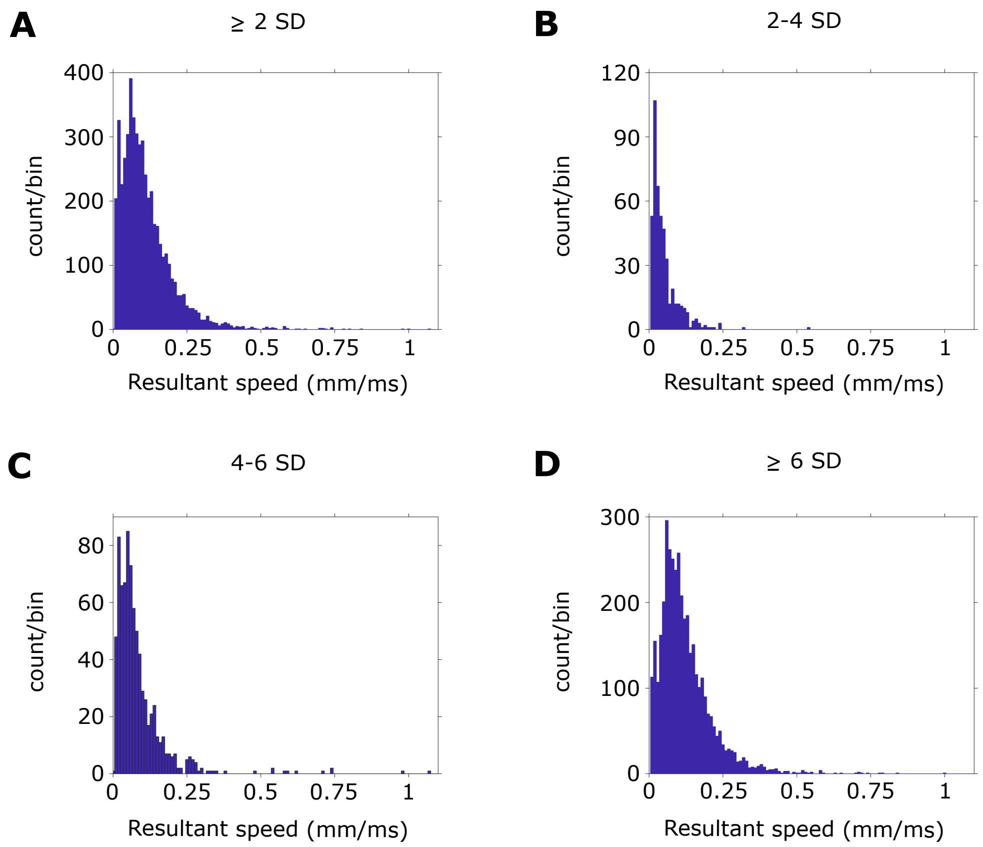


**Supplementary Figure 16** Distribution of resultant speeds.

Distribution of resultant speeds of propagation for ≥ 2 SD (A), 2-4 SD (B), 4-6 SD (C), and ≥ 6 SD (D).

≥ 2 SD: fraction of propagating events with speeds less than 0.2 mm/ms = 0.876
2-4 SD: fraction of propagating events with speeds less than 0.2 mm/ms = 0.983
4-6 SD: fraction of propagating events with speeds less than 0.2 mm/ms = 0.932
≥ 6 SD: fraction of propagating events with speeds less than 0.2 mm/ms = 0.855
